# Myeloid-targeted RNA nanotherapeutics rewire cholesterol metabolism to unleash anti-tumor immunity in glioblastoma

**DOI:** 10.64898/2026.08.03.741800

**Authors:** Jiawei Huo, Hanchen Lin, Yinmeng Li, Shashwat Tripathi, Rafal Chojak, Caylee Silvers, Yirui Peng, Lauren Boland, Jianzhong Zhang, Kathleen McCortney, Rasangi M. Perera, Hinda Najem, Leah K. Billingham, Tzu-Yi Chia, Xiaoyang Chen, Hanxiang Wang, Jingqi Sun, Mark John Siringan, Leon Jing, Amaan Musabji, Harrshavasan Congivaram, Si Wang, Aurora Lopez-Rosas, Priya Kumthekar, Pouya Jamshidi, Atique U. Ahmed, Catalina Lee-Chang, James P. Chandler, Mark W. Youngblood, Adam Sonabend, Matthew C. Tate, Hardik Shah, Edward B. Thorp, Maciej S. Lesniak, Amy B. Heimberger, Jason Miska, Peng Zhang

## Abstract

Tumor-associated myeloid cells (TAMCs) dominate the glioblastoma (GBM) microenvironment and suppress anti-tumor immunity. Here, we identify cholesterol efflux via ABCA1 as a targetable metabolic checkpoint controlling TAMC immunosuppression in GBM. Reprogramming TAMC cholesterol metabolism using TAMC-targeting lipid nanoparticle encapsulating ABCA1 siRNA (ABCA1 LNP) converts TAMCs into potent antigen-presenting cells with enhanced pro-inflammatory activity and antigen-presenting capacity, thereby inducing T cell activation, expansion, and tumor infiltration. Mechanistically, ABCA1 blockade induces cholesterol accumulation in TAMC membranes, promoting lipid raft formation and enhancing MHC-I-mediated antigen presentation. In multiple preclinical GBM models, ABCA1 LNP treatment dramatically induces T cell priming, extends animal survival, and overcomes GBM resistance to radiotherapy and immune checkpoint therapy. This efficacy was well-maintained in stem-like and recurrent GBM models, GBM patient specimens, and a renal cell carcinoma model. Altogether, our work identifies cholesterol efflux as a targetable metabolic vulnerability in TAMCs to overcome therapy resistance in myeloid-rich, immunologically “cold” tumors.

## INTRODUCTION

Glioblastoma (GBM) remains one of the most therapeutically intractable malignancies with an immunosuppressive tumor microenvironment (TME) dominated by tumor-associated myeloid cells (TAMCs)^1–3^. TAMCs are a heterogeneous population of infiltrating myeloid cells, with distinct phenotypes and functions that orchestrate immune dysfunction and constitute a central barrier to effective immunotherapy^4, 5^. Despite their pivotal role in tumor progression and therapeutic resistance, there are no effective tools to selectively target pathways within TAMCs.

Compelling studies have highlighted cholesterol metabolism as a key axis linking cellular metabolism to myeloid immune functionality^6^. In GBM, sustained exposure to tumor-derived lipids drives progressive intracellular lipid accumulation in TAMCs^7^; yet how dysregulated cholesterol handling in these cells shapes the tumor immune microenvironment remains poorly defined. Cholesterol levels dictate membrane organization, receptor clustering, antigen presentation capacity, and inflammatory signaling^8, 9^. Among these regulators, the ATP-binding cassette transporter A1 (ABCA1) is a major mediator of cholesterol efflux^10^. While ABCA1 has been extensively studied in the context of atherosclerosis and other metabolic disorders^11^, its immunological function in tumor-resident myeloid cells is largely unexplored^12^. For example, studies of myeloid cell cholesterol efflux in peripheral tumors are contradictory, with some showing that cholesterol efflux both promotes^13–15^ and inhibits^16^ immunosuppressive myeloid function.

Myeloid-selective modulation of ABCA1 in vivo requires strategies that enable precise, cell-intrinsic gene perturbation. To this end, small interfering RNA (siRNA) offers a robust, reversible, and sequence-specific approach to silencing defined molecular targets^17, 18^, making it particularly suited for mechanistic interrogation of myeloid pathways. Lipid nanoparticles (LNPs) have emerged as a clinically validated delivery platform for RNA therapeutics, offering robust protection, favorable in vivo delivery, and efficient cytosolic release of the payload RNA^19, 20^. Critically, the modular composition of LNPs, along with the rational design of surface functionalization with a targeting ligand, can be tuned to preferentially target defined immune populations.

Here, we engineered LNP-based RNA therapeutics that specifically inhibit ABCA1 in TAMCs (ABCA1 LNP). Building on our antibody-directed immune targeting (ADIT) technology platform^21, 22^, the antibody-functionalized ABCA1 LNP efficiently and specifically delivers ABCA1 siRNA to TAMCs in GBM, leading to robust ABCA1 inhibition and intracellular cholesterol accumulation only in TAMCs. This reprogramming triggers the formation of lipid raft microdomains and enhances major histocompatibility complex class I (MHC-I)-restricted antigen presentation, effectively reprogramming TAMC toward an anti-tumoral pro-inflammatory state that promotes CD8⁺ T cell priming and leads to potent anti-GBM efficacy in therapy-resistant and immunologically “cold” brain tumors.

## RESULTS

### ABCA1 is highly enriched in GBM-associated TAMCs

Pan-cancer dataset^23^ analysis shows a unique enrichment of ABCA1 expression in central nervous system (CNS) malignant tissues, including both high- and low-grade gliomas (GBM and LGG, respectively) (**Fig. 1a, red arrows**). To map the transcriptional landscape of ABCA1, publicly available single-cell RNA sequencing (scRNA-seq) datasets were interrogated from human GBM patients^24^. Within GBM, myeloid cells showed higher ABCA1 expression in a greater fraction of cells, particularly tumor-associated macrophages and monocyte-derived subsets (**Fig. 1b, top panel**). A similar ABCA1 expression pattern was observed in the murine CT-2A GBM model (**Fig. 1b, bottom panel**), validating the utility of the preclinical model for studying myeloid cholesterol metabolism in GBM. Multiplex immunofluorescence analysis of GBM patient specimens (**Table S1**) confirmed the high ABCA1 protein expression with TAMCs across independent patient samples (**Fig. 1c**). In murine models, ABCA1 is enriched in GBM-associated TAMCs relative to bone marrow-derived macrophages (BMDMs) (**Fig. 1d**) or splenic myeloid cells (**Fig. 1e, f**), demonstrating ABCA1 is differentially upregulated in tumors.

**Figure 1.**
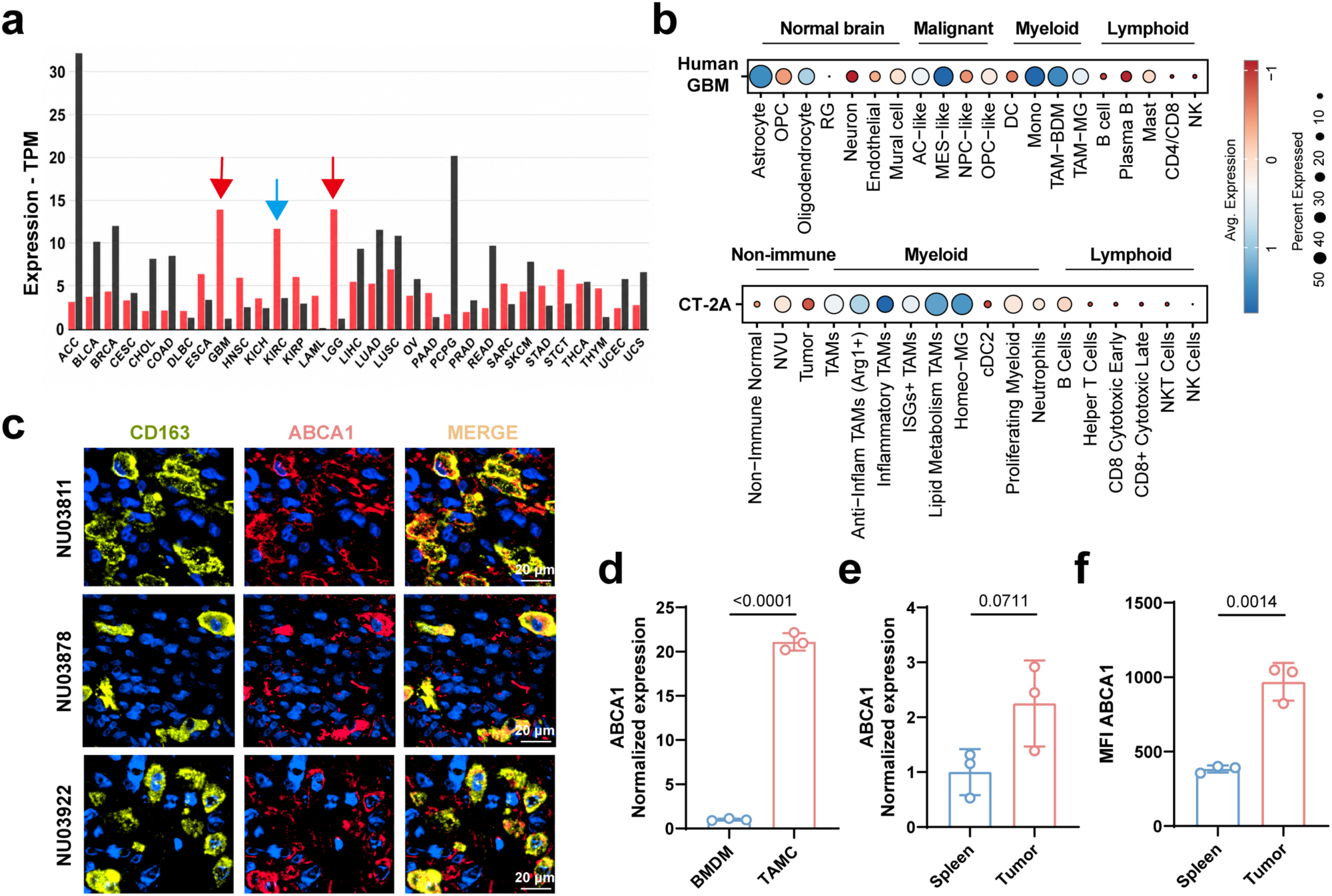
ABCA1 is enriched in GBM-associated TAMCs. **a,** ABCA1 mRNA expression across human cancer types and matched normal tissues (transcripts per million, TPM). Red bars represent cancer tissues and black bars represent matched normal tissues. Red arrows highlight brain cancers with high ABCA1 expression and blue arrow indicates ABCA1 expression in kidney renal clear cell carcinoma (KIRC). **b,** scRNA-seq analysis of ABCA1 transcript levels across major cell populations in human GBM (top) and CT-2A murine GBM (bottom). Dot size represents the percentage of cells expressing ABCA1, and color indicates scaled mean expression. **c,** Multiplex immunofluorescence staining of human GBM specimens with CD163 (yellow), ABCA1 (red), and DAPI (blue), with merged images shown from three independent patient samples (#NU03811, #NU03878, and #NU03922). Scale bars, 20 µm. **d,** Quantitative real-time PCR (qPCR) analysis of ABCA1 mRNA in bone marrow-derived macrophages (BMDMs) versus in vitro-generated TAMCs (n = 3). **e,** RNA-seq analysis of ABCA1 mRNA in splenic myeloid cells and TAMCs freshly isolated from CT-2A tumor-bearing mice (n = 3). **f,** Flow cytometric analysis of ABCA1 expression in myeloid cell populations gated from murine spleen and CT-2A tumors, quantified as mean fluorescence intensity (MFI) (n = 3). Data are presented as the mean ± s.d. Statistical significance for d, e, and f was assessed by unpaired two-tailed Student’s t-test. *p* values are indicated in the figures.

### ABCA1 inhibition disrupts cholesterol efflux and reprograms TAMCs

To interrogate the role of ABCA1 in cholesterol handling by myeloid cells, we encapsulated ABCA1 siRNA into LNPs (ABCA1 LNP, **Fig. 2a**) and delivered them to in vitro TAMCs generated from mouse bone marrow progenitor cells (**Fig. S1)**^21, 22^, which resist conventional transfection^25^. ABCA1 LNP reduced ABCA1 transcripts by over 90% as compared to control LNP encapsulating scrambled siRNA (**Fig. 2b**). This reduction was confirmed at the protein level by flow cytometric analysis (**Fig. 2c**). As a result of the efficient ABCA1 suppression, TAMC total cholesterol levels were robustly increased (**Fig. 2d**), which was further confirmed by liquid chromatography–mass spectrometry (LC-MS) (**Fig. S2a**). Collectively, these data demonstrate that ABCA1 LNP efficiently downregulates ABCA1 in TAMC, driving cholesterol accumulation.

**Figure 2.**
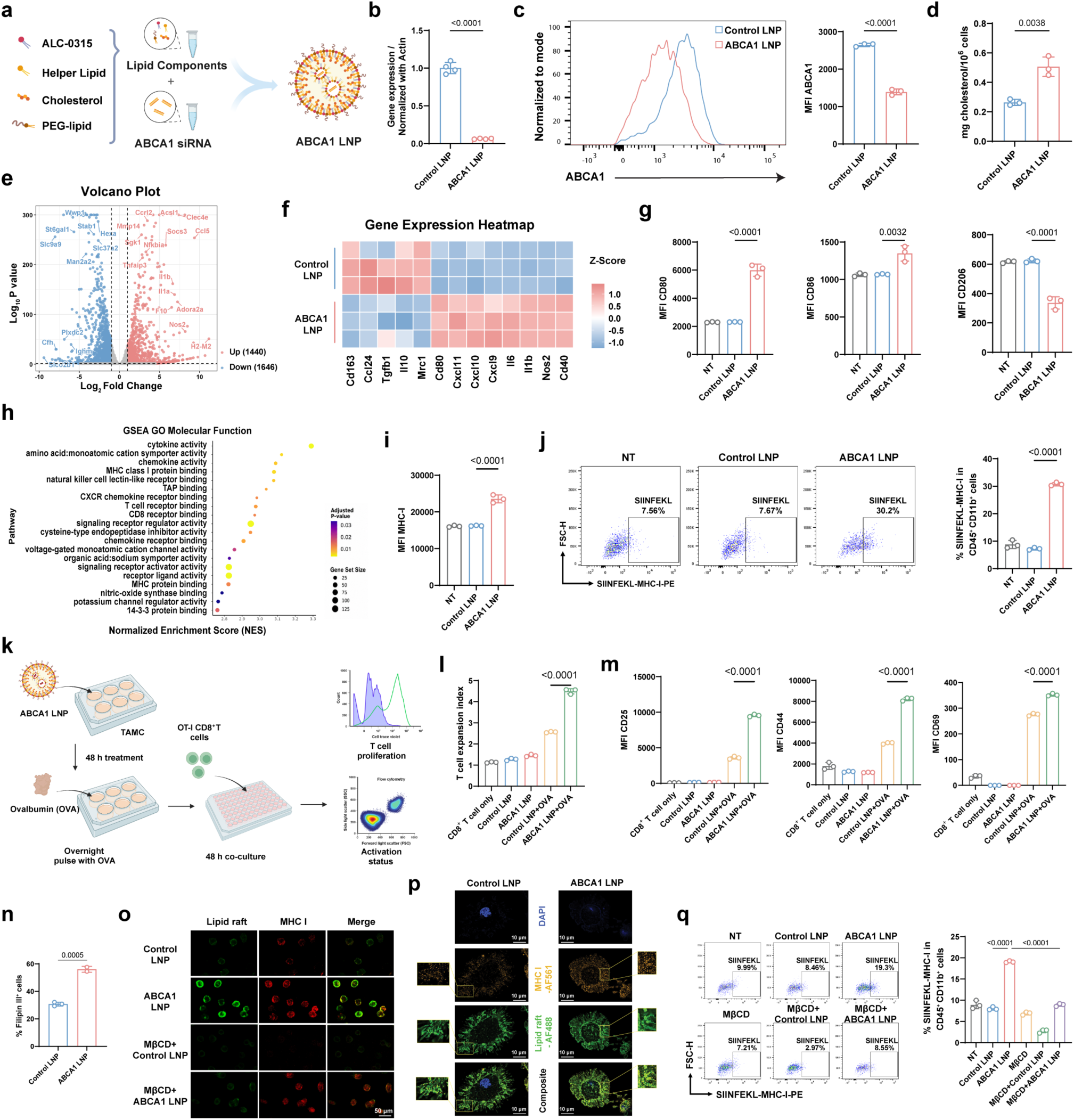
ABCA1 LNP disrupts cholesterol efflux and concentrates membrane cholesterol to promote antigen presentation and TAMC reprogramming. **a,** Schematic of ABCA1 LNP formulation using ionizable lipid (ALC-0315), helper lipid, cholesterol, and PEG-lipid with encapsulated ABCA1 siRNA. **b,** qPCR analysis of ABCA1 mRNA in in vitro-generated TAMCs treated with control LNP (with scrambled siRNA) or ABCA1 LNP at 100 nM siRNA for 24 hours, normalized to β-actin (n = 4). **c,** Representative flow cytometry histograms and quantification of ABCA1 MFI in TAMCs following LNP treatment for 48 hours (n = 3). **d,** Total cholesterol content in TAMCs treated with control LNP or ABCA1 LNP for 48 hours (n = 3). **e,** Volcano plot of bulk RNA-seq comparing TAMCs treated with control LNP versus ABCA1 LNP (n = 3). Up-regulated (1,440) and down-regulated (1,646) genes are indicated in red and blue, respectively (|log2 fold change| > 1, adjusted *p* < 0.05). **f**, Heatmap of selected critical myeloid inflammatory/immunosuppressive genes showing Z-score-normalized expression across treatment conditions (n = 3). **g,** Flow cytometry quantification of MFI for CD80, CD86, and CD206 in TAMCs under no treatment (NT), control LNP, or ABCA1 LNP conditions 48 hours after treatments (n = 3). **h,** GSEA of Gene Ontology molecular function terms using genes ranked by differential expression. **i,** Flow cytometric quantification of MFI for MHC-I in TAMCs based on the indicated conditions (n = 3). **j,** Representative flow cytometry plots and quantification of SIINFEKL-MHC-I complex staining on CD45⁺CD11b⁺ TAMCs 24 hours following co-culture with OVA-expressing CT-2A tumor cells (n = 3). **k,** Schematic of the OT-I CD8⁺ T cell and TAMC co-culture assay. **l,** OT-I CD8⁺ T cell expansion index following co-culture with TAMCs treated with control LNP or ABCA1 LNP, +/- OVA pulsing (n = 3). **m,** Flow cytometry quantification of MFI for CD25, CD44, and CD69 on OT-I CD8⁺ T cells after co-culture with TAMCs under the indicated conditions (n = 3). **n**, Flow cytometry quantification of Filipin III staining in TAMCs treated with control LNP or ABCA1 LNP at 100 nM siRNA for 48 hours (n = 3). **o**, Representative confocal microscopy images of TAMCs treated with control LNP or ABCA1 LNP +/- MβCD and stained for lipid rafts (cholera toxin B subunit–AF488, green) and MHC-I (AF594, red). Merged images are shown in the right column. Scale bar, 50 μm. **p**, Representative high-magnification confocal microscopy images of TAMCs treated with control LNP or ABCA1 LNP, stained with DAPI (blue), MHC-I (AF561, orange), and lipid rafts (cholera toxin B subunit–AF488, green). Composite images are shown in the bottom row. Insets show magnified views of the boxed regions. Scale bars, 10 μm. **q**, Representative flow cytometry plots and quantification of SIINFEKL-MHC-I staining on CD45⁺CD11b⁺ TAMCs following co-culture with OVA-expressing CT-2A tumor cells under NT, control LNP, ABCA1 LNP, MβCD, MβCD + control LNP, and MβCD + ABCA1 LNP conditions (n = 3). Data are presented as the mean ± s.d. Statistical significance for b, c, d, and n was assessed by unpaired two-tailed Student’s t-test; for g, i, j, l, m, and q by one-way ANOVA with Tukey’s multiple-comparisons test. *p* values are indicated in the figures.

Next, we sought to define the global transcriptional impact of ABCA1 knockdown in TAMCs. Bulk RNA sequencing of ABCA1 LNP-treated TAMCs suggested broad transcriptional remodeling relative to control LNP with scrambled-siRNA (**Fig. 2e**). To place these global changes in the context of myeloid polarization states, a heatmap of selected inflammatory and immunoregulatory transcripts was constructed (**Fig. 2f**). This analysis revealed upregulation of pro-inflammatory/myeloid activation markers such as *Nos2*, *Il6*, *Cxcl9*, *Cxcl11*, *Cd80* and downregulation of immunoregulatory/immunosuppressive markers such as *Mrc1*, *Il10*, *Ccl24*, *Cd163* in ABCA1 LNP-treated TAMCs, consistent with a potential shift from immunosuppressive toward a pro-inflammatory state. These signatures were confirmed at the protein level as ABCA1 LNP-treated TAMCs showed increased surface CD80/CD86 and reduced CD206 expression (**Fig. 2g**).

Gene set enrichment analysis (GSEA) also suggested enhancement of antigen presentation machinery, including MHC-I protein binding and T cell receptor binding, along with cytokine and chemokine activity and membrane/trafficking-associated activities, as well as enrichment of membrane-associated and antigen presentation-related functions (**Fig. 2h, Fig. S2b**). Consistent with these findings, MHC-I surface expression rose with ABCA1 LNP treatment (**Fig. 2i**). To determine whether these changes can be translated into enhanced functional antigen presentation, TAMCs were co-cultured with ovalbumin (OVA)-overexpressing CT-2A glioma cells, in which ABCA1 LNP treatment increased SIINFEKL-MHC-I presentation from 8% to 30% (**Fig. 2j**). OVA-pulsed, ABCA1 LNP-treated TAMCs (**Fig. 2k**) drove greater OT-I CD8⁺ T cell expansion (**Fig. 2l, Fig. S2c**) and upregulation of activation markers CD25, CD44, and CD69 (**Fig. 2m**).

As cholesterol-dependent lipid rafts concentrate surface receptors and antigen-presentation machinery for cell-cell interactions^26–29^, we next asked whether ABCA1 inhibition alters lipid raft formation in TAMCs. To assess changes in membrane-associated free cholesterol, TAMCs were stained with Filipin III, a fluorescent probe that binds free membrane cholesterol^30^. Flow cytometric analysis indicated that ABCA1 LNP treatment dramatically increased the percentage of Filipin III⁺ cells (**Fig. 2n, Fig. S2d**), revealing increased membrane-bound cholesterol in TAMCs after ABCA1 blockade. Such membrane-concentrated cholesterol led to markedly increased GM1⁺ lipid rafts and MHC-I in the TAMC plasma membrane, as determined by spatial overlap (**Fig. 2o, p**). This was confirmed by reduced GM1⁺ raft and MHC-I staining after membrane cholesterol depletion with methyl-β-cyclodextrin (MβCD)^31^ (**Fig. 2o**). Critically, ABCA1 LNP-promoted antigen presentation was blunted by MβCD-mediated membrane cholesterol depletion (**Fig. 2q**). Therefore, ABCA1 blockade triggered antigen MHC-I presentation in a cholesterol- and lipid raft-dependent manner, linking ABCA1 LNP-driven cholesterol buildup to membrane raft organization and antigen presentation capability.

### Myeloid-targeted ABCA1 inhibition promotes T cell anti-GBM activity

To enable TAMC-specific ABCA1 blockade in vivo, we developed a TAMC-targeting ABCA1 LNP termed ABCA1 tLNP, which was engineered (**Fig. 3a**) by leveraging an anti-PD-L1-mediated in vivo TAMC targeting mechanism in GBM^21, 22^. ABCA1 tLNP exhibited a spherical morphology and uniform size distribution, with a hydrodynamic diameter of 130.0 ± 0.3 nm, a low polydispersity index (PDI) of 0.12 ± 0.01, and a near-neutral zeta potential of 0.54 ± 0.26 mV (**Fig. 3b**). ABCA1 tLNP had a high targeting efficiency and specificity to GBM-associated TAMCs but not tumor cells or tumor-infiltrating lymphocytes (TILs) (**Fig. 3c**). In the aggressive and therapy-resistant orthotopic CT-2A model^32^, three intracranial doses of ABCA1 tLNP through an implanted cannula^21, 22^ (**Fig. 3d**) significantly extended animal survival, whereas control tLNP encapsulating scrambled siRNA did not show any effect relative to untreated mice (**Fig. 3e**). This benefit was lost in Rag1^-/-^ mice (**Fig. 3f**), implicating adaptive immunity is required for the anti-tumor efficacy of this approach. Flow cytometry profiling (**Fig. 3g, Fig. S3**) confirmed that ABCA1 tLNP induced TAMC-restricted knockdown, with no change in tumor cells or TILs (**Fig. 3h**), and a selective rise in TAMC cholesterol (**Fig. 3i**). Treatment induced increases of TILs, including CD8^+^ and CD4^+^ populations (**Fig. 3j**), with reduced PD-1 and LAG-3 on intratumoral CD8⁺ T cells (**Fig. 3k**). Given the key role of T cells in ABCA1 tLNP therapy, a combination treatment with anti-PD-1 immune checkpoint blockade (αPD-1) was evaluated (**Fig. 3l**). Kaplan-Meier survival analysis revealed a stepwise improvement across treatment groups: median survival (MS) was 21 days for control, 23 days for αPD-1 alone, 34 days for ABCA1 tLNP, and 54.5 days for the combination of αPD-1 and ABCA1 tLNP (**Fig. 3m**), showing a synergistic interaction (Bliss-independence synergy score = +34 days, *p* = 0.0002)^33–35^. Importantly, long-term survivors (LTS) rejected contralateral CT-2A rechallenge (**Fig. 3n**), demonstrating anti-GBM immunologic memory.

**Figure 3.**
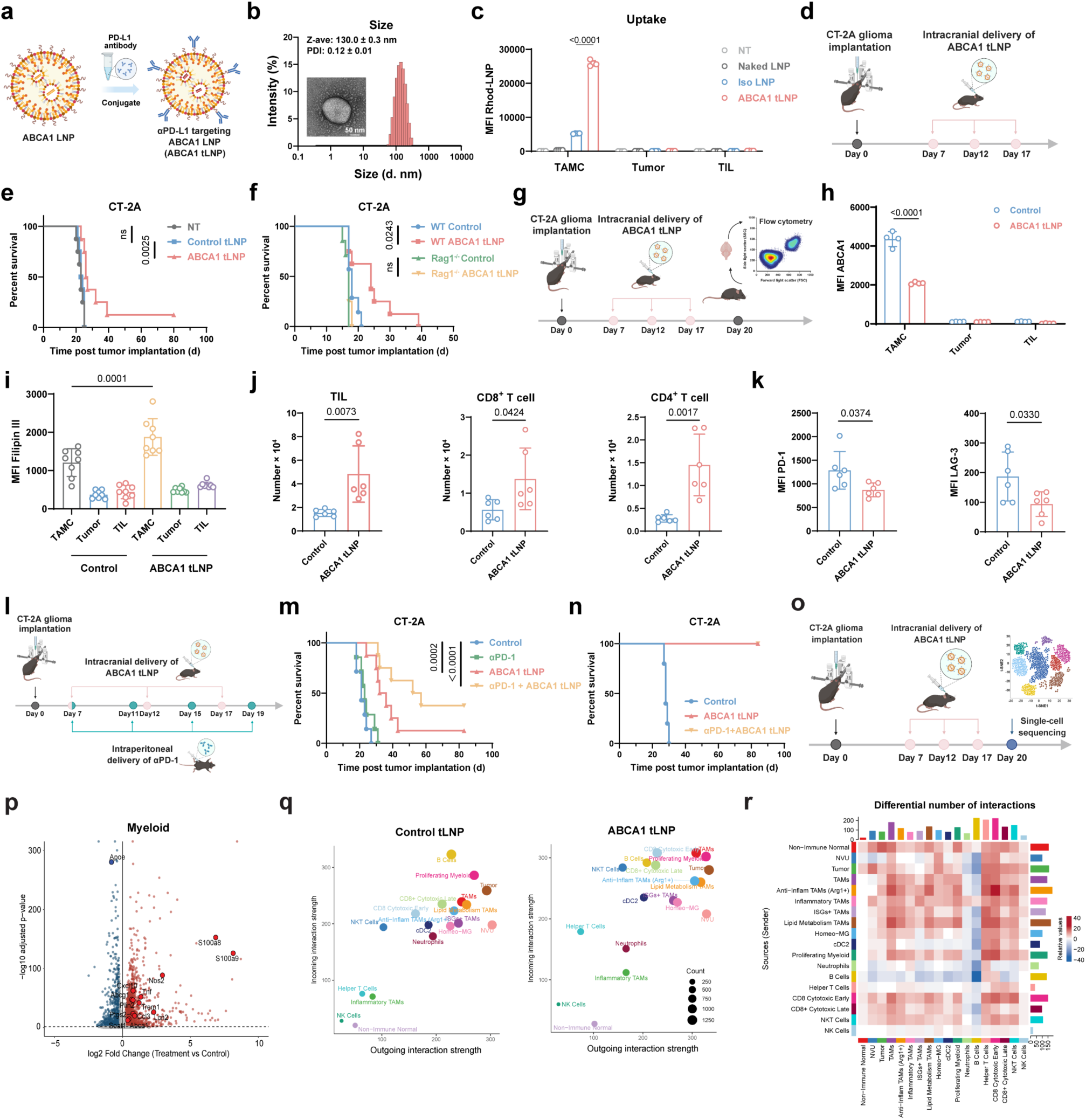
Myeloid-targeted ABCA1 inhibition reshapes transcriptional states and immune signaling networks to promote T cell anti-tumor activity and control GBM progression. **a,** Schematic of ABCA1 tLNP preparation. ABCA1 LNP encapsulating ABCA1 siRNA was conjugated with an anti-PD-L1 antibody (ABCA1 tLNP) for in vivo TAMC targeting in GBM. **b,** Dynamic Light Scattering (DLS) analysis showing the size distribution of ABCA1 tLNP (Z-average 130.0 ± 0.3 nm; PDI 0.12 ± 0.01; n = 3), with a representative TEM image shown as an inset. Scale bar, 50 nm. **c,** Flow cytometric analysis of ex vivo cellular uptake of LNPs. Single-cell suspensions from CT-2A tumors were incubated with the indicated LNPs for 1 hour at 37 °C, and uptake was quantified as MFI by flow cytometry across cell populations (TAMCs, tumor cells, and TILs) under non-treated (NT), naked LNP, isotype-conjugated (IgG2b) LNP (Iso LNP), and ABCA1 tLNP conditions (n = 3). **d,** Schematic of the CT-2A tumor implantation and treatment schedule, including intracranial delivery of ABCA1 tLNP on days 7, 12, and 17 post-tumor implantation. **e,** Kaplan–Meier survival curves of CT-2A-bearing mice treated with non-treated (NT), control tLNP (with scrambled siRNA), or ABCA1 tLNP at 200 μg siRNA/kg (n = 7–8 per group). **f,** Kaplan–Meier survival curves of CT-2A-bearing wild-type (WT) and Rag1⁻/⁻ mice receiving control or ABCA1 tLNP (n = 7–8 per group). **g,** Schematic of the CT-2A tumor immune profiling workflow (for **h-k**) with tumors harvested on day 20 for flow cytometry analysis. **h,** MFI of ABCA1 in TAMCs, tumor cells, and TILs from control and ABCA1 tLNP treatment groups (n = 4). **i,** Flow cytometric quantification of Filipin III staining in TAMCs, tumor cells, and TILs from CT-2A tumors (n = 8). **j,** Numbers of total TILs, CD8⁺ T cells, and CD4⁺ T cells per tumor-bearing brain (n = 6). **k,** MFI of PD-1 and LAG-3 in tumor-infiltrating CD8⁺ T cells (n = 6). **l,** Schematic of CT-2A tumor implantation and the combination treatment schedule, including intracranial delivery of ABCA1 tLNP at 200 μg siRNA/kg on days 7, 12, and 17 and intraperitoneal administration of αPD-1 antibody at 200 μg per mouse on days 7, 11, 15, and 19 post-tumor implantation. **m,** Kaplan–Meier survival curves of CT-2A-bearing mice across mono/combination therapy groups (n = 7–8 per group). **n,** Kaplan–Meier survival curves following contralateral intracranial CT-2A rechallenge of long-term survivors from the ABCA1 tLNP (n = 1) and αPD-1 + ABCA1 tLNP (n = 3) groups, with age-matched mice implanted with CT-2A in parallel as controls (n = 5). **o**, Schematic of experimental design (for **p-r**). CT-2A glioma cells were intracranially implanted on day 0, followed by intracranial administration of ABCA1 tLNP or control tLNP through an implanted cannula at 200 μg siRNA/kg on days 7, 12, and 17. Tumors were harvested on day 20 for scRNA-seq. For each biological replicate, tumors from five mice were pooled. **p**, Volcano plot of differential gene expression in the myeloid compartment. Each dot represents a gene; x-axis indicates log_2_ fold change (treatment vs control tLNP), and y-axis indicates −log_10_ (adjusted *p*-value). Red and blue dots represent significantly up- and down-regulated genes, respectively. Selected representative genes are labeled. **q**, CellChat analysis of outgoing and incoming interaction strength across cell populations in control tLNP- (left) and ABCA1 tLNP-treated (right) groups. Dot size, number of interactions. **r**, Heatmap of differential number of interactions between cell populations (ABCA1 tLNP versus control tLNP). Rows represent signal senders and columns represent signal receivers. The top and right bar plots show the total number of differential interactions per cell type. Color scale indicates the relative change in interaction number (red, increased; blue, decreased in the ABCA1 tLNP-treated group). Data are presented as mean ± s.d. Statistical significance for survival curves (e, f, m, n) was assessed by log-rank (Mantel-Cox) test with Bonferroni correction; c, h, and i by two-way ANOVA with Tukey’s multiple-comparisons test; j and k by unpaired two-tailed Student’s t-test. *p* values are indicated in the figures.

To dissect how ABCA1 targeting reshapes immune signaling networks in GBM, scRNA-seq analysis was performed on orthotopic CT-2A tumors 3 days after the final dose of ABCA1 tLNP (**Fig. 3o**). Compositional analysis^36^ showed expansion of interferon-stimulated gene (ISG)-expressing and pro-inflammatory TAMC subsets at the expense of lipid-metabolic TAMs (**Fig. S4a, b**). Transcriptionally, pro-inflammatory mediators including *Nos2*, *Trem1*, and *S100a8/a9* increased following ABCA1 tLNP treatment (**Fig. 3p**). In T cells, ABCA1 tLNP treatment induced upregulation of genes associated with T cell activation, including Cd44, Runx1, Pim1, Icos, and *Irf4* (**Fig. S4c**). Genes specifically associated with CD4^+^ Th17 cells: *Il17a*, *Il23r*, and *Il17ra* were significantly increased with ABCA1 tLNP treatment (**Fig. S4c**), suggesting Th17 cells may play a role in the anti-tumor response and may be responsible for the increased neutrophils in the LNP treatment group^37^ (**Fig. S4b**). Examination of the tumor compartment revealed a broad upregulation of interferon-stimulated genes, including *Gbp2*, *Cd274 (PD-L1)*, *Stat1*, *Irf1*, and *Oasl2*, indicative of a robust type I/II interferon response (**Fig. S4d**). Cellchat communication analysis^38^ showed strengthened signaling between myeloid subsets and cytotoxic CD8⁺ T cells (**Fig. 3q, r**). Collectively, these data indicate that ABCA1 tLNP drives pro-inflammatory reprogramming of the tumor immune microenvironment, characterized by macrophage activation, induction of the interferon pathway, and augmented immune cell crosstalk.

### Radiation therapy (RT) synergizes with ABCA1-targeted therapy

RT is a key component of the standard of care for GBM patients^39, 40^, and is known to reshape the tumor myeloid compartment^41, 42^. scRNA-seq of irradiated CT-2A tumors showed RT-induced ABCA1 specifically in TAMCs (**Fig. 4a**), and bulk RNA-seq of enriched TAMCs demonstrated the kinetics of ABCA1 upregulation after RT (**Fig. 4b**). Therefore, we next evaluated whether ABCA1 tLNP treatment could synergize with RT to potentiate anti-tumor responses. Given the role of T cells as shown above, a combination with αPD-1 therapy was also tested (**Fig. 4c**). RT + ABCA1 tLNP extended the median survival of CT-2A-bearing animals to 71 days compared with 28 days for RT (*p* = 0.0002). The combination of ABCA1 tLNP with RT + αPD-1 led to effective eradication of CT-2A tumors in approximately 90% of the animals (**Fig. 4d**) (Bliss independence synergy score = +36 days, *p* = 0.0002)^33–35^. Flow cytometric profiling of intratumoral immune populations suggested an increase in the total amounts of TILs, including both CD8⁺ T cells and CD4⁺ T cells, in mice treated with RT + ABCA1 tLNP or RT + αPD-1 + ABCA1 tLNP (**Fig. 4e**). CD8⁺ T cells had reduced expression of exhaustion-associated markers PD-1 and LAG-3 **(Fig. 4f**). In addition, the RT+ABCA1 tLNP (+/- αPD- 1) treatments also induced long-term anti-GBM immune memory in LTS mice, rejecting tumor outgrowth (**Fig. 4g**) and inducing brain-resident T cells with enhanced activation, effector functions, and reduced exhaustion (**Fig. 4h-j, Fig. S5a, b**) in these LTS brains. The crucial role of CD8^+^ T cells in the anti-GBM effectiveness of RT+ABCA1 tLNP combination therapy was further confirmed when the experiment was repeated in CD8-deficient (CD8^-/-^) mice (**Fig. 4k**). Besides CT-2A, in GSC-005, an aggressive, poorly immunogenic glioma stem-like model^43^, ABCA1 tLNP monotherapy modestly extended survival, while the inclusion of RT + αPD-1 effectively cured 3 of 8 animals (37.5%) (**Fig. 4l**). Collectively, these findings suggest a synergy between ABCA1-targeted therapy and RT.

**Figure 4.**
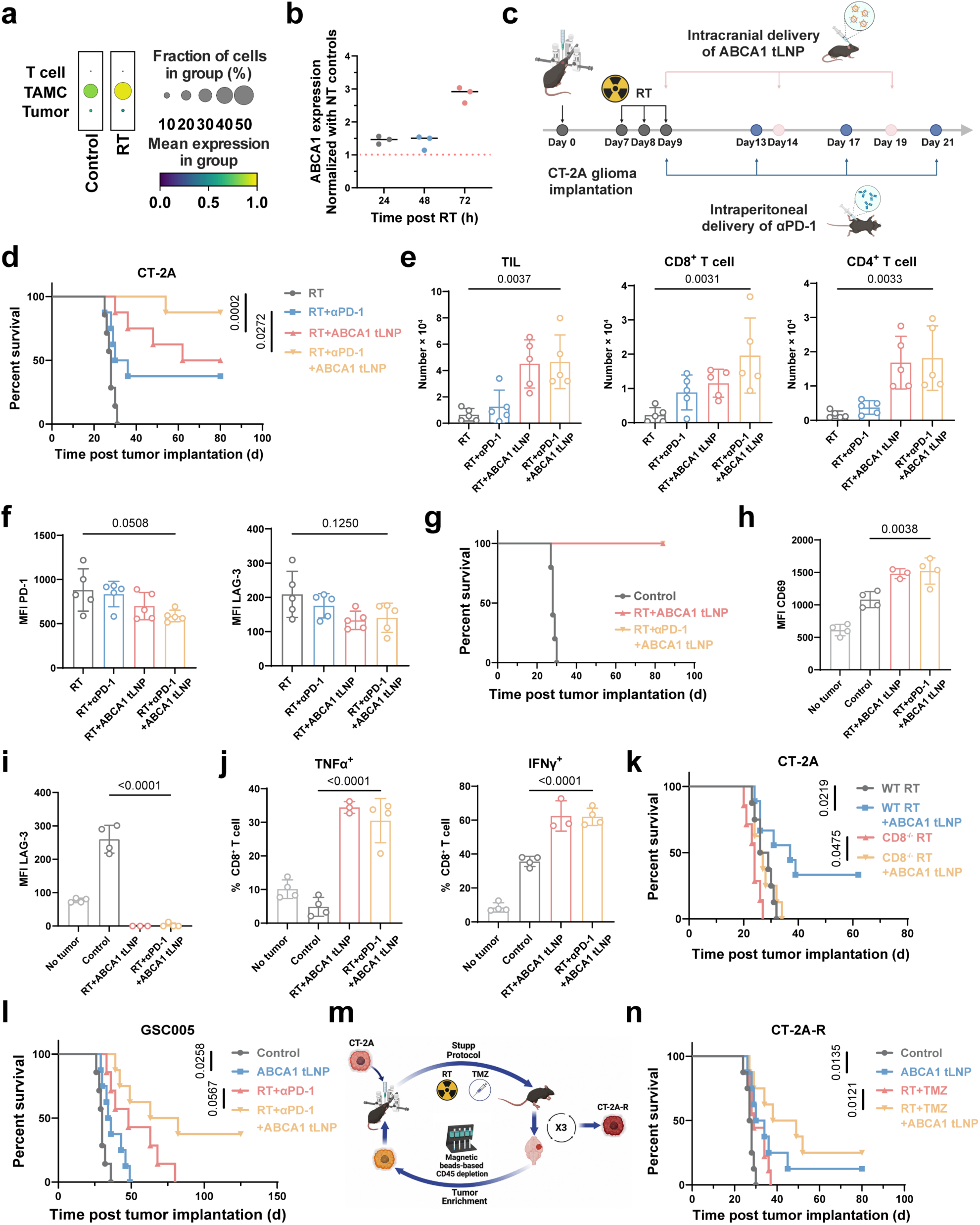
Radiotherapy synergizes with ABCA1 tLNP in multiple preclinical GBM models. **a,** Dot plot of ABCA1 expression in T cells, TAMCs, and tumor cells from control and irradiated (RT) CT-2A tumors. Dot size represents the fraction of cells expressing ABCA1, and color indicates mean expression. **b**, Time course of ABCA1 mRNA expression in TAMCs with magnetic bead enrichment from CT-2A tumors at 24, 48, and 72 hours after RT at 3 × 3 Gy, normalized to the non-irradiated (NT) group (n = 3). **c,** Schematic of orthotopic CT-2A implantation and treatment schedule, including RT on days 7–9, intracranial administration of ABCA1 tLNP at 200 μg siRNA/kg on days 9, 14, and 19, and intraperitoneal administration of αPD-1 antibody at 200 μg per mouse on days 9, 13, 17, and 21. **d,** Kaplan–Meier survival curves of CT-2A-bearing mice implanted intracranially with 50,000 CT-2A cells and treated with RT, RT + αPD-1, RT + ABCA1 tLNP, or RT + αPD-1 + ABCA1 tLNP (n = 7– 8 per group). **e-f**, flow cytometric analysis of the CT-2A tumors after treatments. **e,** Flow cytometric quantification of total TILs, CD8⁺ T cells, and CD4⁺ T cells isolated from tumor-bearing brains under the indicated treatments (n = 5). **f,** MFI of PD-1 and LAG-3 in tumor-infiltrating CD8⁺ T cells under the indicated treatments (n = 5). **g,** Kaplan–Meier survival curves following contralateral intracranial CT-2A rechallenge of long-term survivors previously treated with the indicated regimens. Age-matched tumor-bearing mice implanted with CT-2A in parallel served as controls (Control, n = 5; RT + ABCA1 tLNP, n = 4; RT + αPD-1 + ABCA1 tLNP, n = 7). **h-j**, Flow cytometric analysis of long-term survivors (LTS) brains 80 days after CT-2A tumor rechallenge as compared to control brains. **h,** Flow cytometric quantification of CD69 MFI among brain-infiltrating CD8⁺ T cells from rechallenged mice (n = 4). No tumor indicates age-matched mice without tumor challenge. **i,** Flow cytometric quantification of LAG-3 MFI among brain-infiltrating CD8⁺ T cells from rechallenged mice (n = 4). **j,** Frequencies of TNFα⁺ and IFNγ⁺ cells among brain-infiltrating CD8⁺ T cells from rechallenged mice (n = 4). **k,** Kaplan–Meier survival curves of wild-type (WT) and CD8-deficient (CD8⁻/⁻) mice bearing CT-2A tumors treated with RT or RT + ABCA1 tLNP (n = 7–9 per group). **l,** Kaplan–Meier survival curves of GSC-005–bearing mice treated with non-treated control, ABCA1 tLNP, RT + αPD-1, or RT + αPD-1 + ABCA1 tLNP (n = 7–8 per group). **m,** Schematic of the generation of recurrent CT-2A line (CT-2A-R) cells through three iterative cycles of RT + TMZ treatments in vivo, followed by magnetic bead–based tumor cell enrichment and re-implantation. **n,** Kaplan–Meier survival curves of CT-2A-R–bearing mice implanted intracranially with 1,000 CT-2A-R cells and treated with control, ABCA1 tLNP, RT + TMZ, or RT + TMZ + ABCA1 tLNP (n = 8–9 per group). Due to the high lethality of the CT-2A-R model, a lower implanted cell number was used for treatment feasibility. Data are presented as mean ± s.d. Statistical significance for survival curves (d, g, k, l, n) was assessed by log-rank (Mantel–Cox) test with Bonferroni correction; e, f, h, i, and j by one-way ANOVA with Tukey’s multiple-comparisons test. *p* values are indicated in the figures.

A central challenge of GBM is that tumor recurrence is nearly universal after therapy^40, 44^. To model this setting, we established a recurrent CT-2A line (CT-2A-R) by exposing orthotopic CT-2A tumors to three sequential rounds of the Stupp protocol (fractionated RT + temozolomide (TMZ)-based chemotherapy)^40^, with tumor cell enrichment and re-implantation between cycles (**Fig. 4m**). Compared with parental CT-2A tumors, CT-2A-R tumors displayed markedly reduced sensitivity to RT + TMZ, and intracranial implantation of as few as 500 CT-2A-R cells was sufficient to cause rapid lethality (**Fig. S6**), confirming the therapy-resistant, highly aggressive phenotype of this recurrent model. We then tested whether ABCA1-targeted therapy could enhance tumor response to standard therapy (RT+TMZ) in this recurrent model. Because of the CT-2A-R model’s lethality, the implanted cell number was reduced from 50,000 (CT-2A) to 1,000 (CT-2A-R) to improve treatment feasibility. Although RT+TMZ alone showed marginal anti-tumor effects in CT-2A-R-bearing animals, a combination with ABCA1 tLNP markedly sensitized CT-2A-R tumors to RT+TMZ standard therapy (MS: 29 vs. 43.5 days; *p* = 0.0121) (**Fig. 4n**). Altogether, our data demonstrate the effectiveness of ABCA1 tLNP in multiple preclinical GBM models, including stem-like and recurrent tumors, supporting its translational potential in this refractory disease.

### A human-ready formulation reprograms GBM patient-derived TAMCs

To validate the translatability of ABCA1-targeted therapy, a human-ready ABCA1 tLNP was formulated with anti-human PD-L1 and human ABCA1 siRNA, and applied ex vivo to GBM immune infiltrates dissociated from freshly dissected clinical GBM specimens (**Fig. 5a, Table S1).** As in murine tumors, ABCA1 tLNP uptake was restricted to human TAMCs rather than microglia, tumor cells, or TILs, with selective ABCA1 suppression and increased intracellular cholesterol (**Fig. 5b-d**, **Fig. S7**). Treated TAMCs upregulated inflammatory mediators (IL1B, IL6, CCL2, CXCL10), myeloid activation markers (AIM2, CLEC5A), and lipid remodeling genes (CH25H, SLC27A2, FABP1) (**Fig. 5e, Fig. S8**), with GSEA enrichment of TNFα–NF-κB, interferon-γ and -α, inflammatory response, and IL6-JAK-STAT3 signaling (**Fig. 5f**).

**Figure 5.**
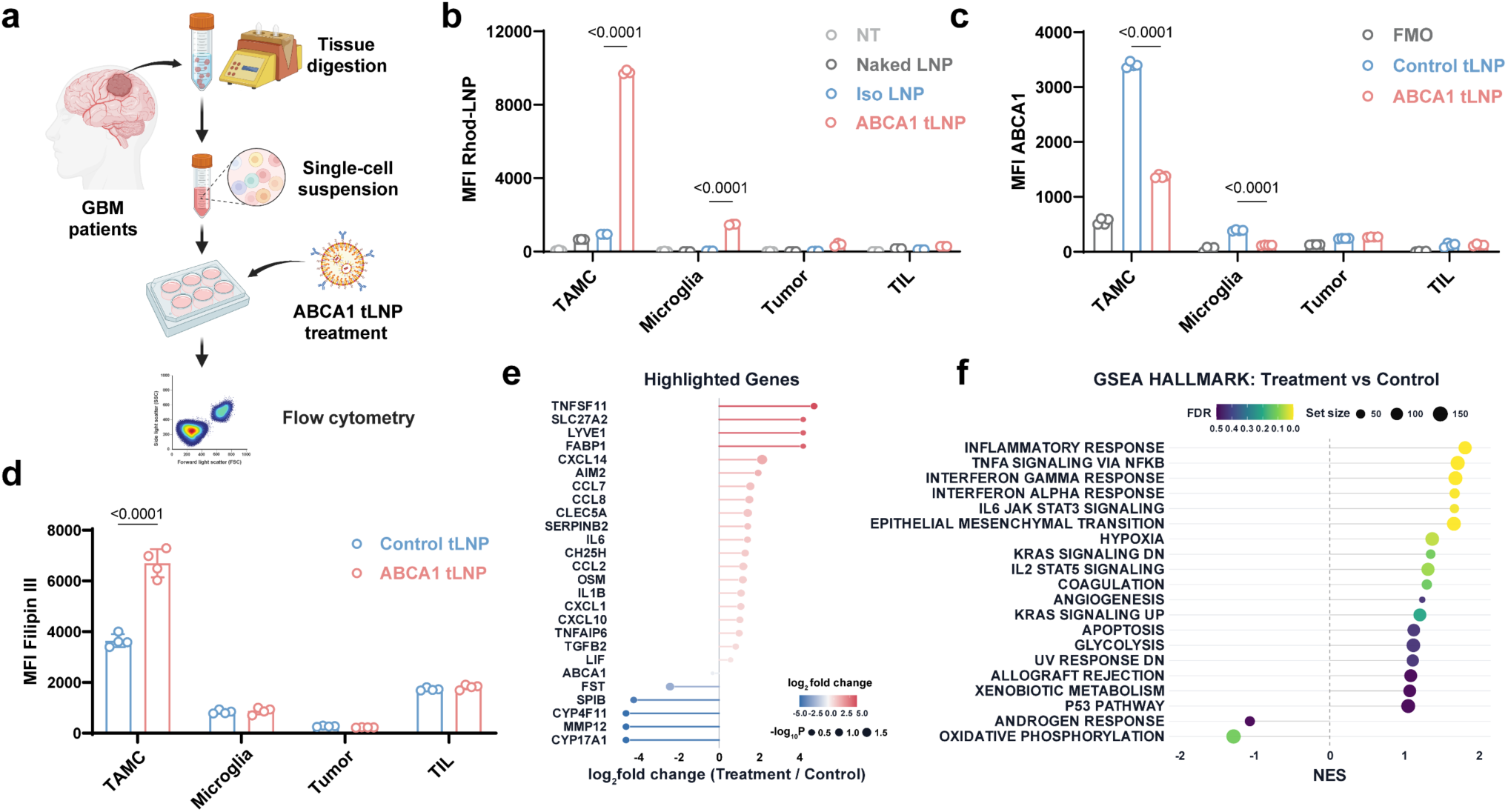
A human-ready ABCA1 tLNP formulation targets and reprograms GBM patient-derived TAMCs. **a,** Schematic of the experimental workflow. Tumor tissues from GBM patients were enzymatically dissociated into single-cell suspensions, followed by ex vivo treatment with designated LNPs and flow cytometric analysis for TAMC targeting, ABCA1 expression, or Filipin III-based cholesterol measurements. **b,** Flow cytometric quantification of LNP uptake (MFI) in TAMCs, microglia, tumor cells, and TILs from human GBM specimen (#NU05064) following 1 hour ex vivo incubation under non-treated (NT), naked LNP, isotype-conjugated (IgG2b) LNP, or ABCA1 tLNP conditions (n = 3). **c,** Flow cytometric quantification of ABCA1 MFI in TAMCs, microglia, tumor cells, and TILs from patient sample #NU05064 (n = 4) following treatment with ABCA1 tLNP or control tLNP (with scrambled siRNA) at 100 nM siRNA for 48 hours. Fluorescence-minus-one (FMO) controls were used to set ABCA1 positivity thresholds and validate staining. **d,** Flow cytometric quantification of Filipin III MFI in TAMCs, microglia, tumor cells, and TILs under control tLNP and ABCA1 tLNP conditions for 48 hours in patient sample #NU04935 (n = 4). **e,** Lollipop plot of selected differentially expressed genes from bulk RNA-seq of CD14⁺ myeloid cells positively selected from dissociated GBM specimens and subsequently treated ex vivo with control tLNP or ABCA1 tLNP for 48 hours (patient sample #NU04887). Bar length represents log_2_ fold change (treatment/control); color indicates log_2_ fold change; dot size represents −log_10_ (*p* value). **f,** GSEA of Hallmark pathways in ABCA1 tLNP-treated versus control tLNP-treated CD14⁺ myeloid cells from patient #NU04887. Dot size represents gene set size; color indicates FDR; x-axis shows normalized enrichment score (NES). Data are presented as mean ± s.d. Statistical significance for b, c, and d was assessed by two-way ANOVA with Tukey’s multiple-comparisons test. *p* values are indicated in the figures.

### ABCA1 tLNP confers anti-tumor efficacy in other ABCA1 overexpressing solid tumors

To determine whether other oncology indications may benefit from ABCA1-targeted therapy, a pan-cancer transcriptomic reference dataset^23^ was queried. Across a spectrum of over 30 cancer types, kidney renal clear cell carcinoma (TCGA: KIRC) displayed a comparably high ABCA1 expression level, alongside GBM and LGG, with elevated expression in malignant versus matched normal tissues (**Fig. 1a, blue arrow**). This analysis, along with the fact that KIRC is well characterized as a myeloid-rich tumor^45^, inspired us to explore the possibility of using ABCA1 tLNP as an immunotherapeutic approach for KIRC.

Single-cell analysis of human renal cell carcinoma (RCC)^46^ strongly indicates that ABCA1 expression was enriched within the TAMC compartment (**Fig. 6a**). Consistent with human data, in a preclinical murine KIRC model, Renca, TAMCs constituted the dominant CD45⁺ immune population (**Fig. 6b**), and these TAMCs highly expressed PD-L1 (**Fig. 6c**), confirming the rationale of using anti-PD-L1-mediated TAMC targeting strategy. Indeed, uptake analysis of rhodamine-labeled ABCA1 tLNP confirmed its preferential accumulation in KIRC-associated TAMCs over tumor cells and TILs (**Fig. 6d**). In TAMCs, ABCA1 knockdown (**Fig. 6e**) dramatically triggered intracellular cholesterol accumulation (**Fig. 6f**), accompanied by upregulation of the T cell-recruiting chemokines *Cxcl9* and *Cxcl11* (**Fig. 6g**). To test the in vivo therapeutic efficacy, BALB/c mice bearing subcutaneous Renca tumors received three intravenous doses of ABCA1 tLNP (**Fig. 6h**), which resulted in significant suppression of tumor growth (**Fig. 6i**) and lower endpoint tumor weight in treated mice (**Fig. 6j**). Immune profiling revealed increased frequencies of both CD8⁺ and CD4⁺ T cells among TILs in ABCA1 tLNP-treated tumors (**Fig. 6k, Fig. S9**), with a concurrent increase in granzyme B-expressing (GzmB⁺) and TNFα⁺ CD8⁺ T cells (**Fig. 6l**). Conversely, expression of the exhaustion-associated markers PD-1 and LAG-3 was reduced in these CD8⁺ T cells (**Fig. 6m**). Altogether, these data support a broad translational potential of ABCA1 tLNP across myeloid-rich, ABCA1-high solid tumors beyond GBM.

**Figure 6.**
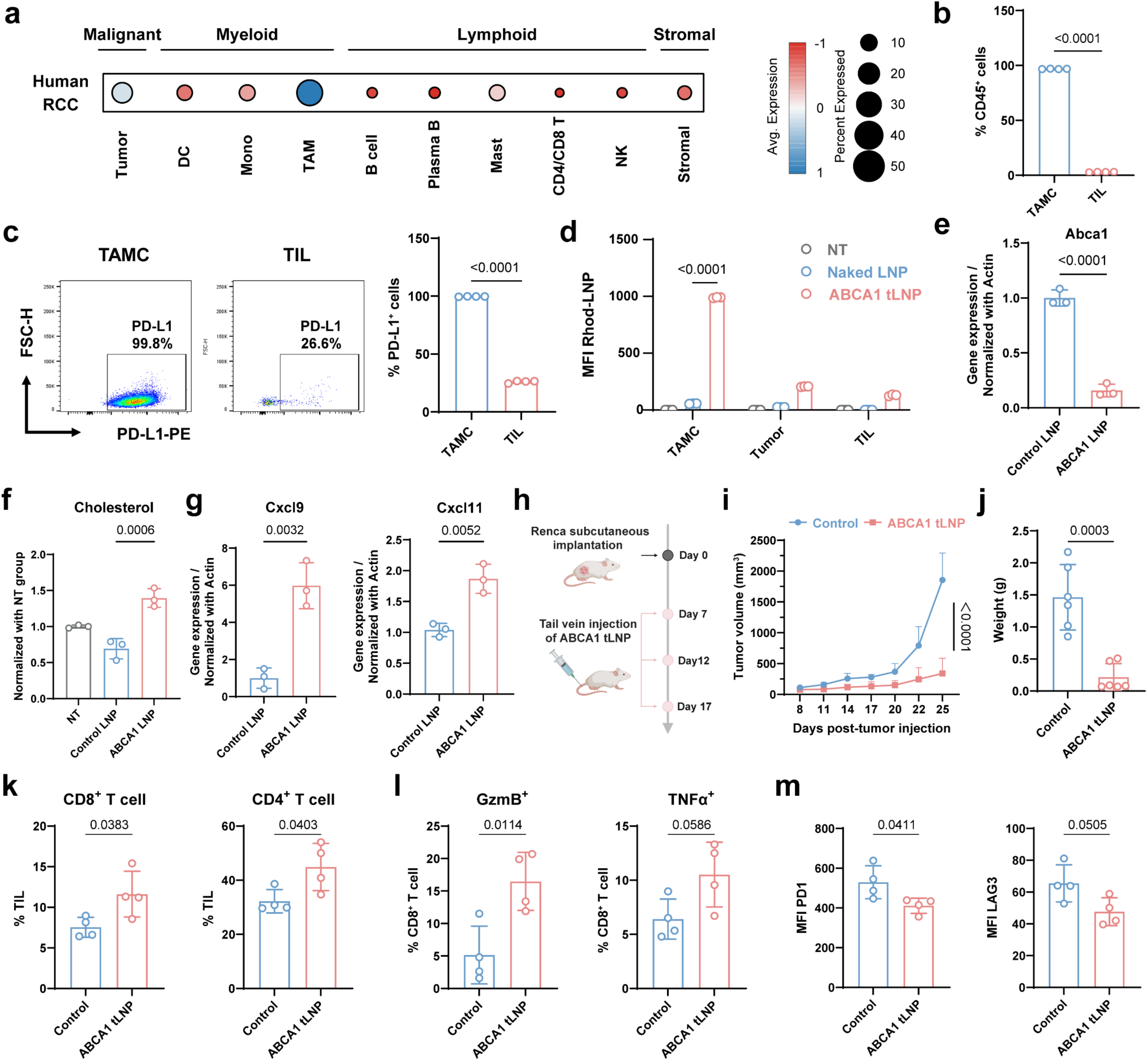
ABCA1 tLNP confers anti-tumor efficacy in ABCA1-overexpressing renal cancers. **a,** Dot-plot analysis of ABCA1 expression across malignant, myeloid, lymphoid and stromal cell populations in human renal cell carcinoma, based on publicly available scRNA-seq data from a cohort of patients with advanced renal cell carcinoma (RCC) (Bi et al.^46^, Single Cell Portal, SCP1288); dot size denotes the percentage of cells expressing ABCA1 and color denotes the average scaled expression level. **b,** Flow cytometric quantification of TAMCs and TILs as a percentage of CD45⁺ cells in Renca renal cancer model (n = 4). **c,** Representative flow cytometry plots and quantification of the frequency of PD-L1⁺ cells among TAMCs and TILs from Renca tumors (n = 4). **d,** MFI of LNP uptake in TAMCs, tumor cells, and TILs under NT, naked LNP, and ABCA1 tLNP conditions for 1 hour at 37 °C (n = 3). **e,** qPCR analysis of Abca1 in Renca-associated TAMCs treated with control LNP or ABCA1 LNP, normalized to the β-actin housekeeping gene and presented relative to the control LNP group (n = 3). **f,** Cholesterol levels in Renca-associated TAMCs treated with NT, control LNP (with scrambled siRNA), or ABCA1 LNP at 100 nM siRNA for 48 hours, normalized to the NT group (n = 3). **g,** qPCR analysis of Cxcl9 and Cxcl11 in Renca-associated TAMCs treated with control LNP or ABCA1 LNP, normalized to the β-actin and presented relative to the control LNP group (n = 3). **h,** Schematic of Renca subcutaneous implantation and tail vein administration schedule for ABCA1 tLNP. **i,** Tumor volume measurement over time following control or ABCA1 tLNP treatment at 800 μg siRNA/kg (n = 6). **j,** Endpoint tumor weight measurement in non-treated control or ABCA1 tLNP-treated mice (n = 6). **k,** Frequencies of CD8⁺ and CD4⁺ T cells within TILs from control and ABCA1 tLNP groups (n = 4). **l,** Frequencies of GzmB⁺ and TNFα⁺ CD8⁺ T cells within TILs from control and ABCA1 tLNP groups (n = 4). **m,** MFI of PD-1 and LAG-3 on tumor-infiltrating CD8⁺ T cells from control and ABCA1 tLNP groups (n = 4). Data are presented as mean ± s.d. Statistical significance for b, c, e, g, and j–m was assessed by unpaired two-tailed Student’s t-test; f by one-way ANOVA with Tukey’s multiple-comparisons test; d and i by two-way ANOVA with Tukey’s multiple-comparisons test. *p* values are indicated in the figures.

## DISCUSSION

In this study, we demonstrate that ABCA1-driven cholesterol efflux functions as a tunable metabolic node that shapes TAMC activation states, immune communication, and responses to RT and immunotherapy-based treatments (as shown in **Table of Contents)**. Rather than a passive consequence of lipid overload^47^, TAMC cholesterol handling emerges as an actionable determinant of immune signaling and therapeutic outcome. A key conceptual finding of this study is the tight coupling between cholesterol efflux and antigen presentation, consistent with studies showing genetic ablation of ABCA1 in macrophages promotes anti-tumor immunity in peripheral tumor models^48^. However, there is still considerable controversy about the role of cholesterol efflux in myeloid cell activities in different cancers^16^, which suggests context-dependent mechanisms are not yet fully elucidated. These data are also consistent with classical evidence that lipid rafts serve as organizing platforms for MHC assembly and T cell receptor engagement^26^.

Single-cell profiling of ABCA1 tLNP–treated tumors extended these findings to the intact tumor ecosystem, revealing coordinated remodeling across myeloid, lymphoid, and malignant compartments. Inflammatory TAMC subsets were enriched over immunoregulatory populations, and the tumor compartment itself displayed an interferon-stimulated gene signature. Cell–cell communication inference further supported tighter functional coupling between activated myeloid subsets and cytotoxic T cells, suggesting that the therapeutic benefit of ABCA1 targeting is driven not by any single cell type in isolation but by network-level restructuring of the GBM immune microenvironment.

The therapeutic outcomes of this reprogramming were most pronounced when ABCA1 targeting was combined with RT. Radiation markedly upregulated ABCA1 in TAMCs, providing a biological rationale for the combination rather than a merely additive effect: RT both induces the target and creates the inflammatory context^49^ in which ABCA1 blockade is most consequential. Consistent with this, the combination of RT, αPD-1, and ABCA1 tLNP produced durable, near-complete responses in orthotopic CT-2A glioma and retained activity in therapy-resistant GSC-005 and recurrent CT-2A-R models, where standard RT + TMZ had lost efficacy. The requirement for CD8⁺ T cell involvement, demonstrated by loss of benefit in CD8-knockout hosts and protection against contralateral rechallenge in long-term survivors, indicates that ABCA1 tLNP therapy generates immunologic memory. Given that recurrent GBM remains nearly uniformly lethal and is the setting in which most clinical trials are initiated, efficacy in the CT-2A-R model is of translational interest.

In summary, our study identifies ABCA1 not merely as a cholesterol transporter but as a central regulator coordinating antigen presentation, chemokine signaling, and immune activation within the GBM microenvironment. Beyond the specific biology of ABCA1, the myeloid-directed LNPs offer a generalizable strategy broadly applicable to immunologically “cold”, myeloid-rich, and immunotherapy-resistant cancers.

## METHODS

### Animals

C57BL/6J (RRID:IMSR_JAX:000664), OT-I (RRID:IMSR_JAX:003831), and BALB/c (RRID:IMSR_JAX:000651) mice were obtained from The Jackson Laboratory and housed in a pathogen-free barrier facility at Northwestern University. Animals were kept on a 14 hours light/10 hours dark cycle at 22°C with relative humidity maintained between 30% and 70%. Male and female mice aged 6–8 weeks were randomly allocated to control or treatment groups and co-housed across experimental conditions to minimize cage effects. All animal procedures were performed under protocols approved by the Institutional Animal Care and Use Committee (IACUC) of Northwestern University.

### Human samples

Human GBM specimens were acquired from the Northwestern University Nervous System Tumor Bank (NSTB) under protocols approved by the Northwestern University Institutional Review Board (IRB No. STU00095863 and STU00202003). Patients undergoing neurosurgical resection for nervous system tumors who met the study inclusion criteria were enrolled in the tumor tissue bank after providing written informed consent at the time of surgical consent. All procedures were performed in accordance with the ethical principles of the U.S. Common Rule. Each case was reviewed by a board-certified neuropathologist, and tumors were classified according to the 2021 WHO classification of central nervous system tumors. For this study, fresh specimens from patients with newly diagnosed GBM (NU04887, NU05064, and NU04935) were used for tissue dissociation and downstream analyses, and specimens from three additional patients (NU03811, NU03878, and NU03922) were used for multiplex immunofluorescence.

### Reagents

Lipid components were obtained from Avanti Polar Lipids, including 1,2-distearoyl-sn-glycero-3-phosphocholine (DSPC), [(6-hydroxybutyl)azanediyl]dihexane-6,1-diyl bis(2-hexyldecanoate) (ALC-0315), 1,2-dimyristoyl-rac-glycero-3-methoxypolyethylene glycol-2000 (DMG-PEG2000), 1,2-distearoyl-sn-glycero-3-phosphoethanolamine-N-[maleimide(polyethylene glycol)-2000] (DSPE-PEG2000-maleimide), and L-α-Phosphatidylethanolamine-N-(lissamine rhodamine B sulfonyl) (Rhod-PE). Cholesterol and ABCA1 small interfering RNA (siRNA) were sourced from Sigma-Aldrich. Temozolomide (TMZ) was purchased from Cayman Chemical. InVivoMAb anti-mouse PD-1 (clone RMP1-14), anti-mouse PD-L1 (clone 10F.9G2), and anti-human PD-L1 (clone 29E.2A3) antibodies were obtained from Bio X Cell. All reagents were used as received. The siRNA sequences used in this study are listed in **Table S2**.

### Cell lines

The CT-2A glioma cell line used in this study was kindly provided by Dr. Tom Seyfried (Boston College). The glioma stem cell line GSC-005 was kindly provided by Dr. Inder M Verma (Salk Institute). The Renca murine renal cell carcinoma line was purchased from the American Type Culture Collection (ATCC). The CT-2A cell line overexpressing ovalbumin (CT-2A-OVA) was generated as previously described^22^. To generate treatment-resistant CT-2A-R, CT-2A tumors were orthotopically implanted in C57BL/6J mice and treated from day 7 with brain-focused fractionated RT (3 Gy × 3) plus intraperitoneal TMZ (50 mg/kg × 5). Tumors were enzymatically dissociated, CD45⁺ cells were depleted with magnetic beads, and remaining tumor cells were reimplanted. This harvest-selection-reimplantation cycle was repeated three times.

CT-2A and CT-2A-R cells were maintained in Dulbecco’s Modified Eagle Medium (DMEM; Corning) supplemented with 10% fetal bovine serum (FBS; HyClone) and 1% penicillin-streptomycin (Invitrogen). CT-2A-OVA cells were maintained in the same medium with addition of G418 (Sigma) at 200 μg/mL. The Renca cells were cultured in RPMI-1640 (Corning) supplemented with 10% FBS and 1% penicillin-streptomycin. The GSC-005 cell line was cultured in ReNcell NSC medium (Millipore) supplemented with N2 (Invitrogen), B27 without vitamin A (Invitrogen), 20 ng/mL FGF-2 (R&D Systems), and 20 ng/mL EGF (R&D Systems). All cell lines were maintained at 37°C in a humidified atmosphere containing 5% CO₂ and were routinely tested for mycoplasma contamination.

### Formulation and characterization of ABCA1 LNP and ABCA1 tLNP

ABCA1 LNPs were formulated by ethanol injection. Briefly, ALC-0315, cholesterol, DSPC, and DMG-PEG2000 were dissolved in chloroform at a molar ratio of 50.5:37:10:2.5. For fluorescently labeled LNPs, Rhod-PE was incorporated into the lipid mixture at 0.5% (molar ratio). The organic solvent was evaporated under a gentle nitrogen stream, and the resulting lipid film was subsequently dried under vacuum for 4 hours to remove the residual chloroform. The dried lipids were reconstituted in anhydrous ethanol and rapidly mixed with ABCA1 siRNA in citrate buffer (10 mM, pH 4.0) at an aqueous-to-ethanol volume ratio of 3: 1. The mixture was then dialyzed against Dulbecco’s phosphate-buffered saline (DPBS) to exchange buffer and adjust the final formulation to physiological pH (7.4). Control LNPs were prepared in parallel using a non-targeting scrambled siRNA under identical conditions.

For antibody conjugation to generate targeted LNPs (tLNPs), LNPs were formulated using the same procedure except that DMG-PEG2000 was replaced with DSPE-PEG2000-maleimide to enable thiol–maleimide coupling on the particle surface. Anti-mouse PD-L1 antibody (clone 10F.9G2, Bio X Cell) was thiolated using 2-iminothiolane (Traut’s reagent, Sigma) in PBS (pH 8.0) containing 4 mM EDTA at a 2-iminothiolane-to-antibody molar ratio of 10: 1. After incubation at room temperature for 1 hour, excess 2-iminothiolane was removed using Amicon Ultra centrifugal filters (10 kDa MWCO; Millipore). The thiolated antibody was then conjugated to freshly prepared LNPs at a ratio of 80 μg antibody per mg total lipid with overnight incubation at 4°C under gentle rotation, and then purified by centrifugation. The physicochemical characteristics of the formulations were assessed using a Zetasizer Nano ZSP (Malvern) to measure hydrodynamic diameter and zeta potential for antibody-conjugated LNPs. Particle morphology was analyzed by transmission electron microscopy (TEM) after negative staining.

### In vitro generation of TAMCs

Bone marrow progenitor cells were flushed from the tibiae and femurs of C57BL/6J mice as previously described^21^. Cells were seeded into 6-well plates at 1 × 10⁶ cells per well in complete RPMI medium [RPMI-1640 (Corning) containing 10% FBS, 1% penicillin-streptomycin, 1% HEPES (Sigma), 1% non-essential amino acids (Gibco), 1% sodium pyruvate (Corning), and 0.1% 2-mercaptoethanol (Gibco) supplemented with 40 ng/mL M-CSF (PeproTech)]. After 72 hours, the medium was exchanged with a 1:1 mixture of fresh complete RPMI and CT-2A-conditioned medium. Conditioned medium was collected from CT-2A cultures plated at 2 × 10⁶ cells per dish and harvested after 72 hours. Cells were cultured for additional 72 hours, and acquisition of the immunosuppressive TAMC phenotype was confirmed by flow cytometry.

### Isolation and activation of T cells

Splenocytes were harvested from C57BL/6J mice, and T cells were enriched using the MagniSort Mouse T Cell Enrichment Kit (Invitrogen). Enriched T cells were maintained in complete RPMI medium and activated with CD3/CD28-coated Dynabeads Mouse T-Activator (Gibco). Recombinant murine IL-2 (PeproTech) was added to a final concentration of 50 U/mL. For OT-I experiments, CD8⁺ T cells were isolated from the spleens of OT-I TCR-transgenic mice and used directly without CD3/CD28 bead activation.

### In vitro cholesterol measurement, antigen presentation, T cell proliferation assay

Free and total cholesterol were quantified in TAMCs following 48 hours of treatment with control LNP or ABCA1 LNP using a cholesterol quantification assay kit (Abcam) and normalized to cell number. To assess antigen presentation, TAMCs were treated with control LNP or ABCA1 LNP for 48 hours and subsequently co-cultured with OVA-expressing CT-2A cells (CT-2A-OVA) at a 1:1 ratio for 24 hours. Surface presentation of the OVA-derived peptide SIINFEKL bound to MHC class I (H-2Kᵇ) on TAMCs was detected by flow cytometry using an anti-mouse H-2Kᵇ/SIINFEKL antibody (BioLegend). For OT-I T cell proliferation assays, TAMCs were treated with ABCA1 LNP for 48 hours and then pulsed overnight with 10 μg/mL OVA protein (Sigma). After thorough washing to remove unbound antigen, OVA-pulsed TAMCs were co-cultured with CellTrace-labeled OT-I CD8⁺ T cells at a TAMC-to-T cell ratio of 1:2 for 48 hours. T cell proliferation was quantified by flow cytometric analysis based on CellTrace dye dilution and activation status was evaluated by surface staining for CD25, CD44, and CD69.

### Quantitative PCR

Total RNA was extracted using the RNeasy Mini Kit (Qiagen, Hilden, Germany). RNA concentration and purity (A260/A280) were assessed with a NanoDrop spectrophotometer (Thermo Fisher). Equal amounts of RNA were reverse transcribed into complementary DNA (cDNA) using the iScript cDNA Synthesis Kit (Bio-Rad). Quantitative real-time PCR (qPCR) was performed on a CFX96 (Bio-Rad) using the SYBR Green master mix (Bio-Rad). Relative gene expression was calculated using the 2^−ΔΔCt method and normalized to β-actin. Primer sequences are listed in **Table S3**.

### Immunofluorescence (IF) staining and confocal imaging

TAMCs were plated on FluoroDish (FD35-100) and treated with control LNP or ABCA1 LNP for 48 hours. For lipid raft labeling, live cells were labeled with 1 μg/mL Alexa Fluor 488-conjugated cholera toxin B (CTB-AF488; Invitrogen) for 20 min at 4°C, fixed with 4% paraformaldehyde for 10 min at room temperature, washed three times with PBS, and blocked with 2.5% BSA/PBS for 1 hour. For MHC-I staining, samples were incubated overnight at 4°C with anti-MHC class I antibody (Invitrogen; 1:300), then for 1 hour at room temperature in the dark with goat anti-mouse secondary antibody (Invitrogen); nuclei were counterstained with DAPI. For MβCD rescue, TAMCs were pretreated with 5 mM methyl-β-cyclodextrin (Sigma-Aldrich) for 30 min at 37°C before CTB staining. Images were acquired with identical settings at Northwestern University’s Center for Advanced Microscopy and Nikon Imaging Center on a Nikon AXR microscope with a galvano unidirectional scanner, NSPARC detector, and Plan Apo λD 60× oil objective (NA 1.42; OFN25 DIC). Two-dimensional Richardson-Lucy deconvolution used NIS-Elements v5.22.00 (Nikon).

### Bulk RNA-seq analysis

Gene-level counts were generated from BAM files using featureCounts from the Rsubread package with GENCODE v49 annotation. In the two-sample workflow, the GTF annotation was harmonized to match the BAM contig naming convention and re-exported before counting. In the multi-sample workflow, an exon-plus-intron SAF annotation was derived from GENCODE v49 and used for counting. Primary alignments were retained, and multimapping and multi-overlapping reads were excluded. Count matrices were analyzed in edgeR, normalized using TMM, and filtered to remove lowly expressed genes before downstream analysis. Differential expression analysis was performed using edgeR::exactTest with a fixed BCV of 0.40. In the single-comparison workflow, nominal P values were used for visualization. Exploratory pathway analysis included rank-based GSEA.

### Orthotopic mouse GBM models, cannula implantation, and radiotherapy

Orthotopic GBM models were generated in equal numbers of male and female 6–8-week-old C57BL/6J mice by stereotactic intracranial glioma implantation as described^21^. Unless indicated, mice received 5 × 10⁴ CT-2A, 1 × 10⁴ of GSC-005, or 1 × 10^3^ of CT-2A-R cells. For experiments requiring repeated intracranial delivery of therapeutic agents, a guide cannula (Protech International) was surgically implanted prior to tumor implantation to enable serial intracranial administration^21^. RT began on day 7 and comprised 3 Gy/day for three days (9 Gy total) using a Gammacell 40 Exactor irradiator (Best Theratronix). Procedures and monitoring were approved by the Northwestern University IACUC. Mice were euthanized by CO₂ inhalation followed by cervical dislocation at predefined endpoints: persistent loss of righting reflex, limb stiffness, seizures, circling, motor weakness, inability to access food or water, or body-condition score <2.

### Isolation of glioma-infiltrating immune cells

Tumor-bearing mice were euthanized under approved IACUC protocols and transcardially perfused with 5 mL of ice-cold DPBS to clear circulating blood cells. Whole tumor-bearing brains were harvested and mechanically dissociated in Hanks’ Balanced Salt Solution (HBSS; Gibco) using a Potter-Elvehjem tissue grinder fitted with a PTFE pestle. The resulting cell suspension was separated on a discontinuous Percoll (GE Healthcare) density gradient (30% over 70%) to isolate mononuclear cells from myelin and cellular debris. Mononuclear cells were collected from the 30%/70% interphase, washed twice with HBSS, and resuspended in complete RPMI medium for downstream phenotyping and ex vivo functional assays.

### In vivo ABCA1 tLNP treatment combined with radiotherapy

C57BL/6J mice bearing orthotopic CT-2A tumors (5 × 10⁴ cells per mouse) were randomized into treatment groups on day 7 post-implantation. Focal cranial RT was delivered as described above. Anti-PD-1 antibody (clone RMP1-14, Bio X Cell) was administered intraperitoneally at 200 μg per mouse on days 9, 13, 17, and 21 post-tumor implantation. Control or ABCA1 tLNP was delivered intracranially through the implanted guide cannula at 200 μg/kg siRNA on days 9, 14, and 19 post-implantation using a neuros syringe (Hamilton). Survival was monitored according to predefined endpoint criteria. Long-term survivors (LTS) were defined as mice surviving beyond 80 days after tumor implantation, and these mice were subsequently subjected to contralateral intracranial tumor rechallenge or immune profiling. For immune memory studies, age-matched tumor-bearing or control healthy mice from parallel cohorts were euthanized at the indicated time points, and brains were harvested for downstream flow cytometry and histological analyses.

### Flow cytometry and immunophenotyping

All antibodies used for flow cytometric analyses were obtained from BioLegend and applied at a dilution of 1:200, unless otherwise noted. To reduce nonspecific binding, Fc receptors were blocked using anti-CD16/32 antibodies prior to staining single-cell suspensions. Cell viability was determined using the Fixable Viability Dye eFluor780 (Fisher), and dead cells were excluded from downstream analyses. CD45 BV510 and CD11b BV711 were used to determine the immune and myeloid compartments. For assessment of cholesterol levels, Filipin III staining was performed following completion of surface marker labeling. Intracellular cytokine production was measured by stimulating cells for 4 hours with a cell stimulation cocktail plus protein transport inhibitors (Fisher). Cells were then fixed and permeabilized using the Foxp3 fixation/permeabilization (Invitrogen) protocol. Data were acquired on a BD FACSymphony flow cytometer using FACSDiva software and analyzed with FlowJo. Detailed gating strategies are provided in the Supplementary Figures, and antibody information is listed in **Table S4**.

### Single-cell RNA sequencing analysis

C57BL/6 mice with orthotopic CT-2A tumors received intracranial treatment (200 μg/kg) through a cannula on days 7, 12, and 17; tumors were collected on day 20. Each biological replicate pooled tumors from five mice. Tumors were dissociated with the Adult Brain Dissociation Kit and gentleMACS Dissociator (Miltenyi Biotec). After anti-CD16/32 Fc blockade, immune cells were enriched with CD45 MicroBeads (Miltenyi Biotec), and CD45⁺ and CD45⁻ cells were mixed 7:3. scRNA processing was conducted using prior established standard protocols^36, 50, 51^. Post-library preparation cells were sequenced using the Illumina NovaSeq through Northwestern Sequencing Core (NUSeqCore). Raw sequencing files were aligned to mm10 reference using Cell Ranger (v3.1.0). The Seurat R Package using the scRNA-seq Seurat10x genomic workflow was used for all analyses unless noted otherwise^52^. Doublets were removed using scDblFinder with 0.15 threshold. Cells were filtered using a percent mitochondrial DNA threshold of 5% and a UMI range of 100 to 7500. Cells were then subjected to Log Normalize, Scale Data, and PCA functions. The FindClusters and FindMarkers functions were utilized for clustering and marker identification. Non-linear dimensional reduction techniques were applied to visual data in UMAP plot format. The Harmony algorithm was used to regress batch effects allowing for early stop and max iterations of 10^53^. Immune cell clusters were annotated using existing references against the differentially expressed genes found using FindAllMarkers^36^. Ligand-receptor prediction analysis was conducted using CellChat v2 using standard thresholds and functions and results were visualized using netVisual_heatmap, netAnalysis_signalingRole_scatter and netVisual_aggregate.

### Multiplex Immunofluorescent Staining

Formalin-fixed, paraffin-embedded (FFPE) human glioblastoma tissue specimens were sectioned at 5 μm thickness and mounted onto glass slides. Sections were deparaffinized using BOND Dewax solution. Heat-induced epitope retrieval was performed for 20 minutes using either BOND epitope retrieval solution (pH 6) or EDTA buffer (pH 9), followed by endogenous peroxidase quenching and protein blocking. Primary antibodies were diluted in 1× Opal Antibody Diluent/Block solution and paired with the indicated Opal fluorophores: DAPI, GFAP, CD31, CD163, CD11c, P2RY12, CD8, CD4, and ABCA1 with corresponding Opal fluorophore. Multiplex staining was performed using a sequential tyramide signal amplification workflow. In each staining cycle, slides underwent antigen retrieval, blocking, primary antibody incubation, HRP-conjugated secondary antibody labeling, and Opal fluorophore deposition. Following completion of each round, antibody complexes were stripped using heat-mediated retrieval before proceeding to the next marker. After all targets were labeled, sections were counterstained with spectral DAPI and mounted using ProLong Diamond Antifade Mountant.

### Validation of human-ready ABCA1 tLNP formulation using clinical specimens

Fresh human GBM tissue was immediately rinsed in cold PBS, minced, and dissociated with the Adult Brain Dissociation Kit and gentleMACS Dissociator (Miltenyi Biotec), including debris and red-blood-cell removal, then resuspended in complete RPMI-1640. Human-ready ABCA1 tLNP was prepared as above using human ABCA1 siRNA and anti-human PD-L1 (clone 29E.2A3, Bio X Cell). To assess targeting, patient-derived tumor and immune cells were incubated with Rhod-PE-labeled LNPs for 1 hour at 37°C, and uptake MFI was measured by flow cytometry. For functional studies, single-cell suspensions were treated ex vivo with control or ABCA1 tLNP (100 nM siRNA) for 48 hours, then analyzed by flow cytometry for ABCA1 and Filipin III. For bulk RNA-seq, CD14⁺ TAMCs were selected with MojoSort Human CD14 Nanobeads (BioLegend), treated ex vivo with control or ABCA1 tLNP for 48 hours, and harvested for RNA extraction.

### Renca subcutaneous model

Renca cells at a concentration of 1 × 10⁶ in 100 μL PBS were implanted subcutaneously in BALB/c mice (6-8 weeks old). Tumor growth was monitored using caliper measurements, and tumor volumes were calculated over time. ABCA1 tLNPs were administered intravenously via tail vein injection at 800 μg/kg siRNA per injection on days 7, 12, and 17 after tumor implantation. Mice were euthanized when tumors exceeded 1000 mm³, and tumors were collected and weighed at the experimental endpoint. For immune profiling, tumors were harvested three days after the final ABCA1 tLNP dose. Excised tumors were rinsed with cold PBS, minced into 2–4 mm pieces, and dissociated using the High Activity Tumor Tissue Enzymatic Digestion Kit (Mouse) (RWD Life Science) in combination with a gentleMACS Dissociator (Miltenyi Biotec). The resulting cell suspension was passed through a pre-wetted 70 μm cell strainer, and erythrocytes were removed by red blood cell lysis. Cells were then washed, resuspended in complete RPMI-1640 medium, and used for flow cytometric analysis.

### Statistical analysis

Analyses used GraphPad Prism v11 unless noted. Two-group comparisons used two-tailed Student’s t-tests; comparisons of more than two groups used one-way ANOVA with Tukey’s test; longitudinal multigroup comparisons used two-way ANOVA with Tukey’s correction. Data are mean ± s.d. Survival was analyzed by Kaplan-Meier curves and log-rank (Mantel-Cox) tests, with Bonferroni correction where appropriate. Treatment synergy analyses were evaluated under the Bliss independence and highest-single-agent (HSA) models and performed in Python using the lifelines package with custom scripts implementing the Bliss-independence framework for survival data^33–35^.

## AUTHOR INFORMATION

### Author Contributions

P.Z., J.M., and J.H. conceived and designed the study. J.H., H.L., Y.L., C.S., Y.P., L, B., J.Z., L.K.B., T.C., X.C., H.W., J.S., M.J.S., L.J., A.M., and S.W. performed experiments and acquired data. S.T. and R.C. provided assistance with bioinformatic analysis. H.N. and H.C. provided assistance with multiplex immunofluorescence staining and imaging analysis. R.M.P. and H.S. performed LC-MS and data analysis. A.L.-R. performed animal breeding for the study. K.M., P.J., J.P.C., M.W.Y., A.S., and M.C.T. provided supports with human glioblastoma specimens and associated clinical annotation. J.H., H.L., J.M., and P.Z. analyzed data and interpreted results. P.K., A.U.A., C.L-C., E.T., A.B.H. and M.S.L. provided critical scientific input and clinical perspective. J.H., J.M., and P.Z. wrote the manuscript with input from all authors. All authors reviewed the manuscript and approved the final version.

### Conflicts of Interest

A provisional patent application pertaining to the work presented in this manuscript was filed by Northwestern University. The remaining authors declare no competing interests relevant to the specific content of this manuscript.

## ACKNOWLEDGEMENTS

We would like to thank the Northwestern Nervous System Tumor Bank for providing samples from patients with glioblastoma. We would also like to thank Northwestern University Flow Cytometry Core Facility, Analytical bioNanoTechnology Core Facility, and NUSeq Core Facility, which are supported in part by the Northwestern University Cancer Center Support Grant (CCSG; NCI P30CA060553). This work was supported by National Institutes of Health (NIH)/National Cancer Institute (NCI) grants (R37CA266487 to P.Z., R01CA279686 to J.M.), Northwestern Brain Tumor SPORE Developmental Research Program (P50CA221747 to P.Z. and J.M.), and the Collaborative Pilot Award by Center for Human Immunobiology at Northwestern University to P.Z. and J.M. L.B. was also supported by NIH/NCI training grant T32CA268935 at Northwestern University and the Brinson Medical Research Fellowship through The Brinson Foundation.

---

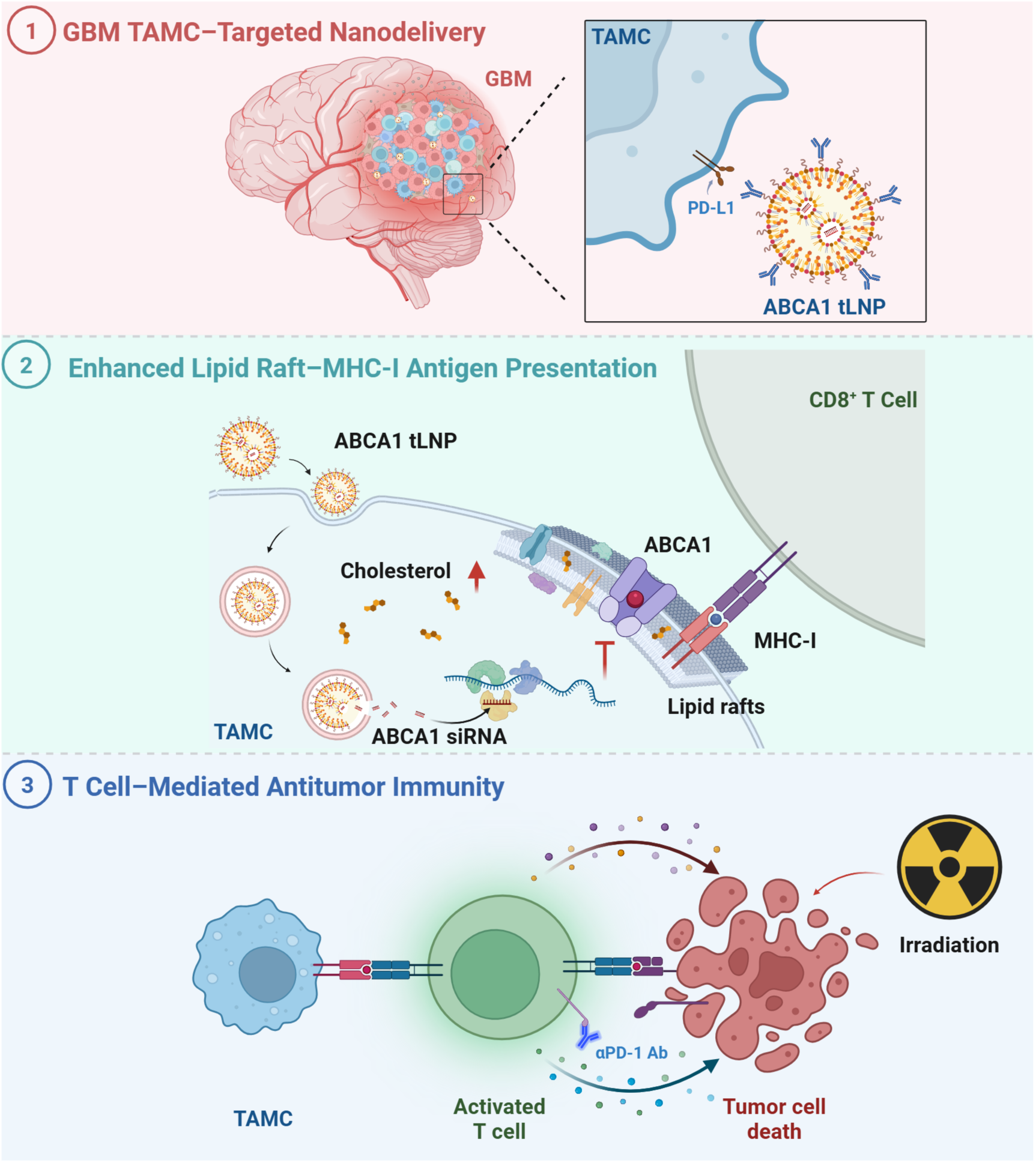
Table of Contents: Myeloid-targeted RNA nanotherapeutics rewire cholesterol metabolism to unleash anti-tumor immunity in glioblastoma. **(1)** Antibody-functionalized lipid nanoparticles encapsulating ABCA1 siRNA (ABCA1 tLNP) efficiently and specifically deliver ABCA1 siRNA to GBM-associated myeloid cells, but not tumor cells or tumor-infiltrating lymphocytes. **(2)** ABCA1 inhibition disrupts cholesterol efflux and drives cholesterol accumulation within TAMC membranes, triggering the formation of lipid raft microdomains and enhancing MHC-I-restricted antigen presentation. **(3)** Reprogrammed TAMCs promote CD8⁺ T cell priming and tumor cell killing, an effect potentiated by αPD-1 blockade and radiotherapy. Created with BioRender.

