## Supplementary Information for "Myeloid-targeted RNA nanotherapeutics rewire cholesterol metabolism to unleash anti-tumor immunity in glioblastoma"

**Figure S1. In vitro TAMC generation and flow cytometry gating strategy.**

**Figure S2. Additional in vitro characterization of ABCA1 LNP-treated TAMCs.**

**Figure S3. Flow cytometry gating strategy for mouse CT-2A tumor samples.**

**Figure S4. Single-cell RNA sequencing of CT-2A tumors following ABCA1 tLNP treatment.**

**Figure S5. Flow cytometry plots of TNFα and IFNγ expression in $CD8^+$ T cells.**

**Figure S6. Kaplan–Meier survival characterization of the CT-2A-R model.**

**Figure S7. Flow cytometry gating strategy for human glioblastoma samples.**

**Figure S8. Volcano plot of differentially expressed genes from bulk RNA-seq comparing ABCA1 tLNP-treated versus control tLNP-treated $CD14^+$ myeloid cells from GBM patient sample.**

**Figure S9. Flow cytometry gating strategy for Renca tumor samples.**

**Table S1. Characteristics of GBM patient samples.**

**Table S2. siRNA sequences used in this study.**

**Table S3. qPCR primer sequences used in this study.**

**Table S4. Antibodies used for flow cytometry and immunophenotyping.**

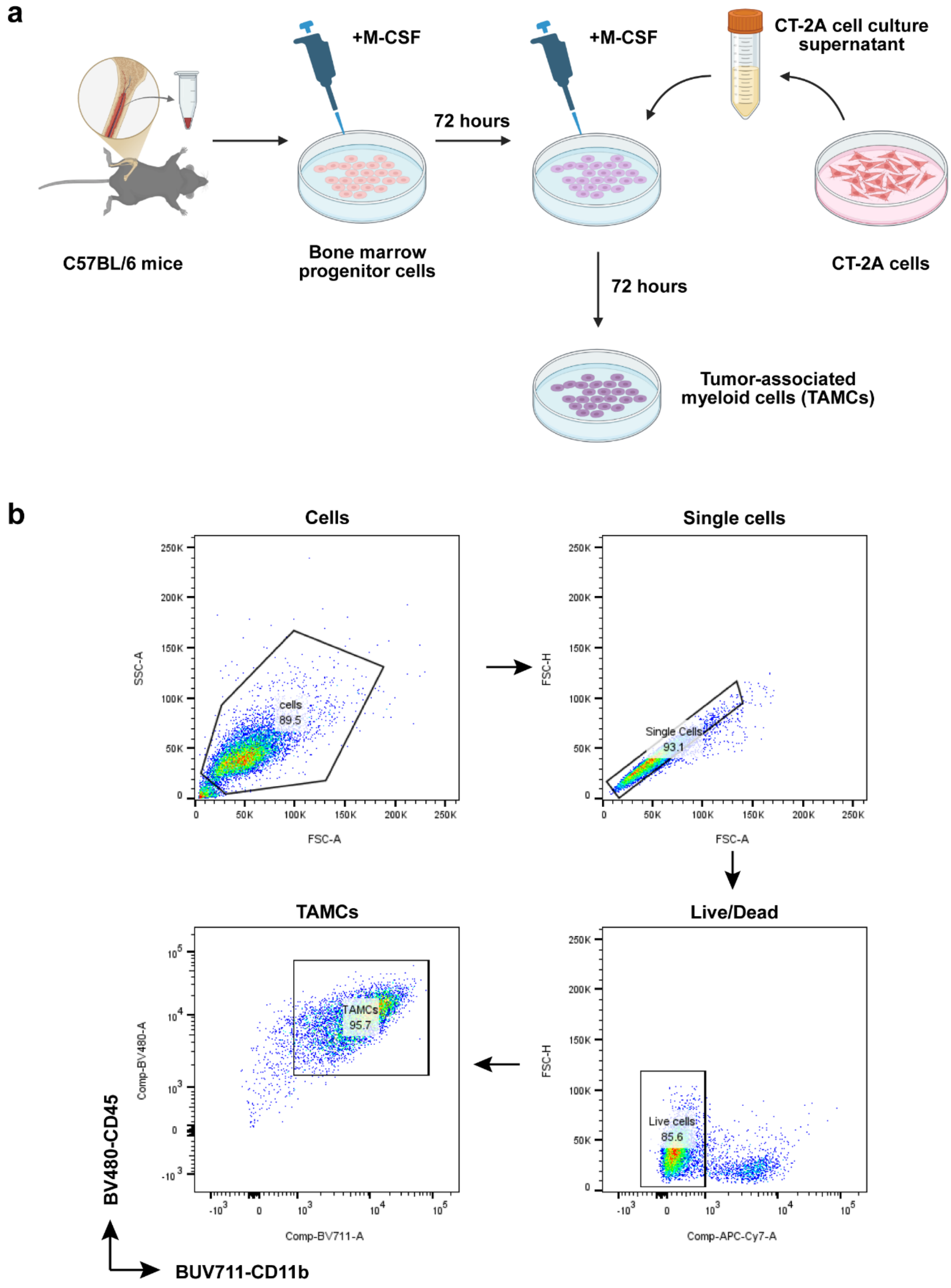


**Figure S1. In vitro TAMC generation and flow cytometry gating strategy. a**, Schematic representation of in vitro TAMC generation. Bone marrow progenitor cells isolated from C57BL/6 mice were differentiated with M-CSF and conditioned with CT-2A cell culture supernatant to generate TAMCs. **b**, Flow cytometry gating strategy for in vitro generated TAMCs. TAMCs were identified following sequential gating on cells, single cells, and live cells.

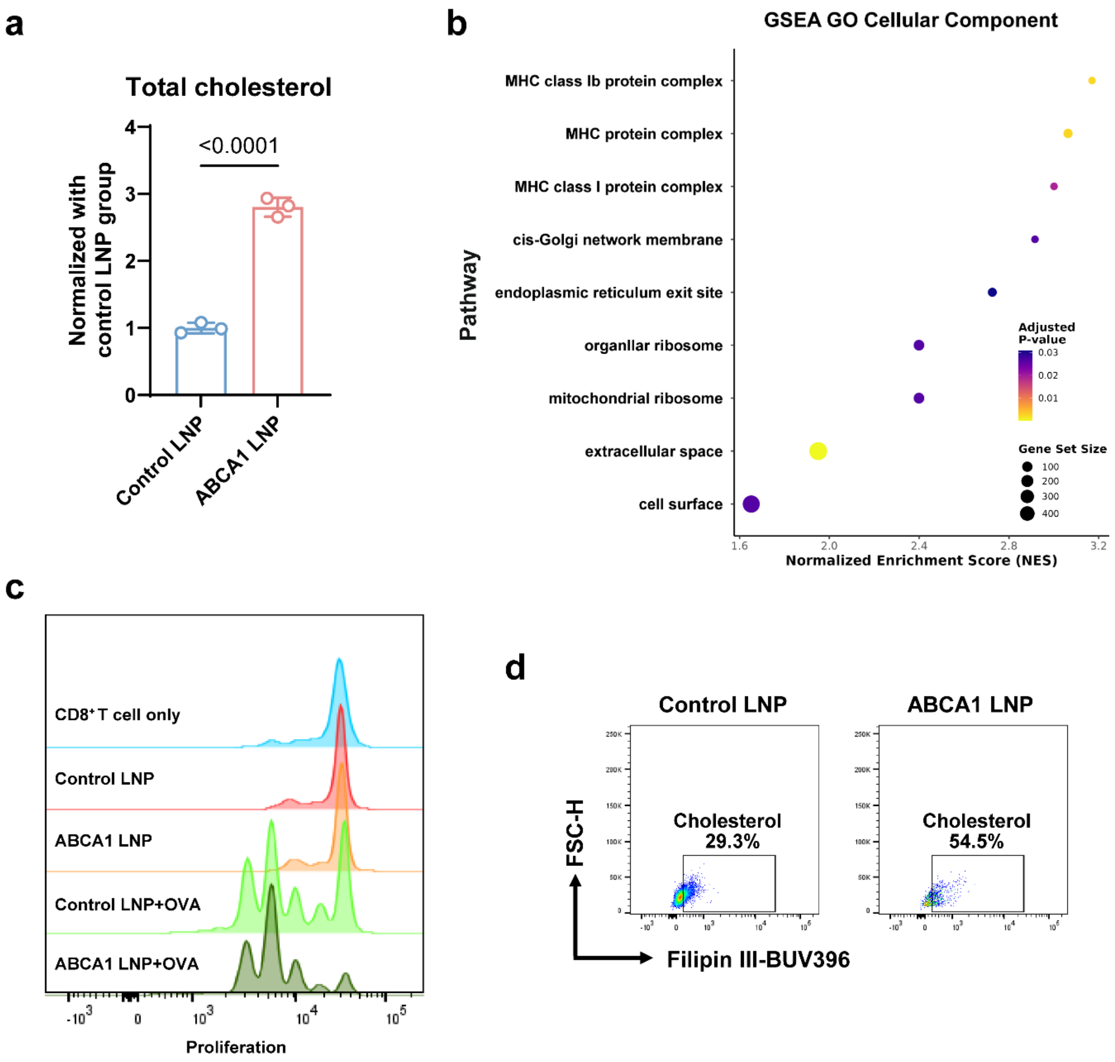


**Figure S2. Additional in vitro characterization of ABCA1 LNP-treated TAMCs. a**, LC-MS quantification of total cholesterol in TAMCs. TAMCs were treated with control LNP and ABCA1 LNP for 48 hours. Cholesterol levels are normalized to the control LNP group. **b**, GSEA of Gene Ontology (GO) cellular component terms using genes ranked by differential expression. Dot size represents gene set size; color indicates adjusted *p* value; x-axis shows the normalized enrichment score (NES) for each pathway. **c**, Flow cytometric analysis of OT-I $CD8^+$ T cell proliferation. Treatment conditions included $CD8^+$ T cells alone, control LNP, ABCA1 LNP, control LNP + OVA, and ABCA1 LNP + OVA. Representative of n = 3. **d**, Representative flow cytometry plots of Filipin III staining in TAMCs treated with control LNP or ABCA1 LNP at 100 nM siRNA for 48 hours (n = 3), with the percentage of Filipin III-positive (cholesterol high) events shown in each gate. Data are presented as mean ± s.d. Statistical significance for a was assessed by unpaired two-tailed Student's t-test, and the *p* value is indicated in the figure.

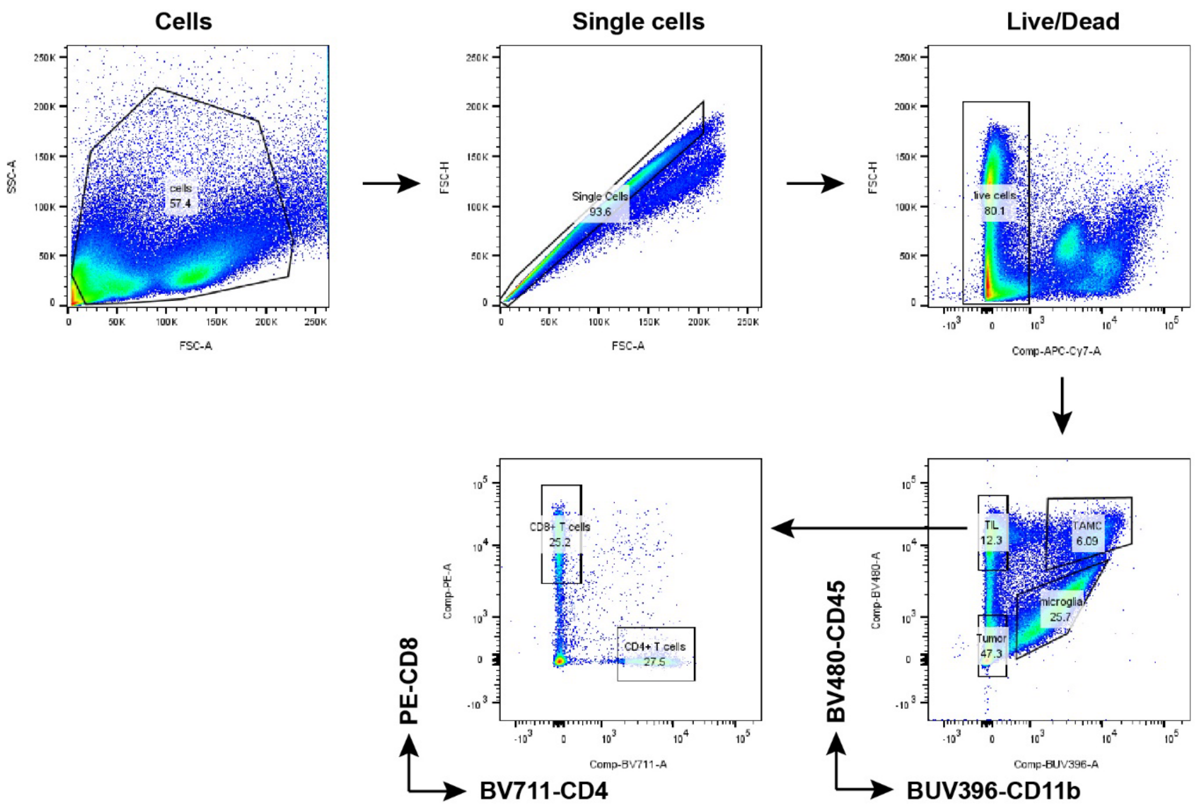


**Figure S3. Flow cytometry gating strategy for mouse CT-2A tumor samples.** Cells were sequentially gated on total cells (FSC-A vs SSC-A), singlets (FSC-A vs FSC-H), and live cells. Populations were then resolved by CD45 and CD11b expression into TAMCs ($CD45^{high}$ $CD11b^{+}$), microglia ($CD45^{int}$ $CD11b^{+}$), tumor-infiltrating lymphocytes (TILs; $CD45^{+}$ $CD11b^{-}$), and tumor cells ($CD45^{-}$). TILs were further separated into $CD8^{+}$ and $CD4^{+}$ T cell populations based on CD8 and CD4 expressions.

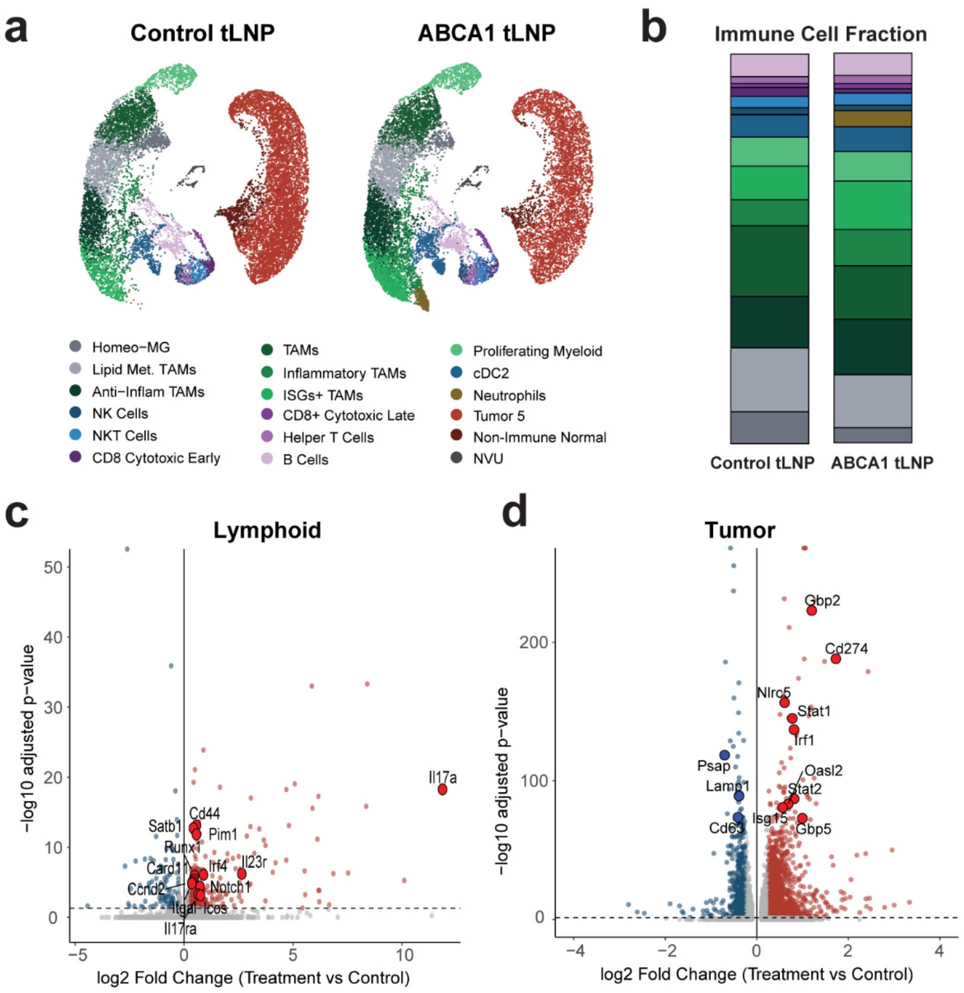


**Figure S4. Single-cell RNA sequencing of CT-2A tumors following ABCA1 tLNP treatment. a**, UMAP projection of the major cell populations identified from scRNA-seq of control tLNP and ABCA1 tLNP-treated tumors, annotated by cluster as previously described[36]. **b**, Composition analysis of the indicated immune and non-immune cell populations in control tLNP and ABCA1 tLNP groups. Color coding matches the cluster labels in **a**. **c, d**, Volcano plots of differential gene expression between ABCA1 tLNP-treated and control tLNP groups in the lymphoid (**c**) and tumor (**d**) compartments. Each dot represents a gene; x-axis indicates $\log_2$ fold change (treatment vs control LNP), and y-axis indicates $-\log_{10}$ (adjusted p-value). Red and blue dots represent significantly up- and down-regulated genes, respectively. Selected representative genes are labeled.

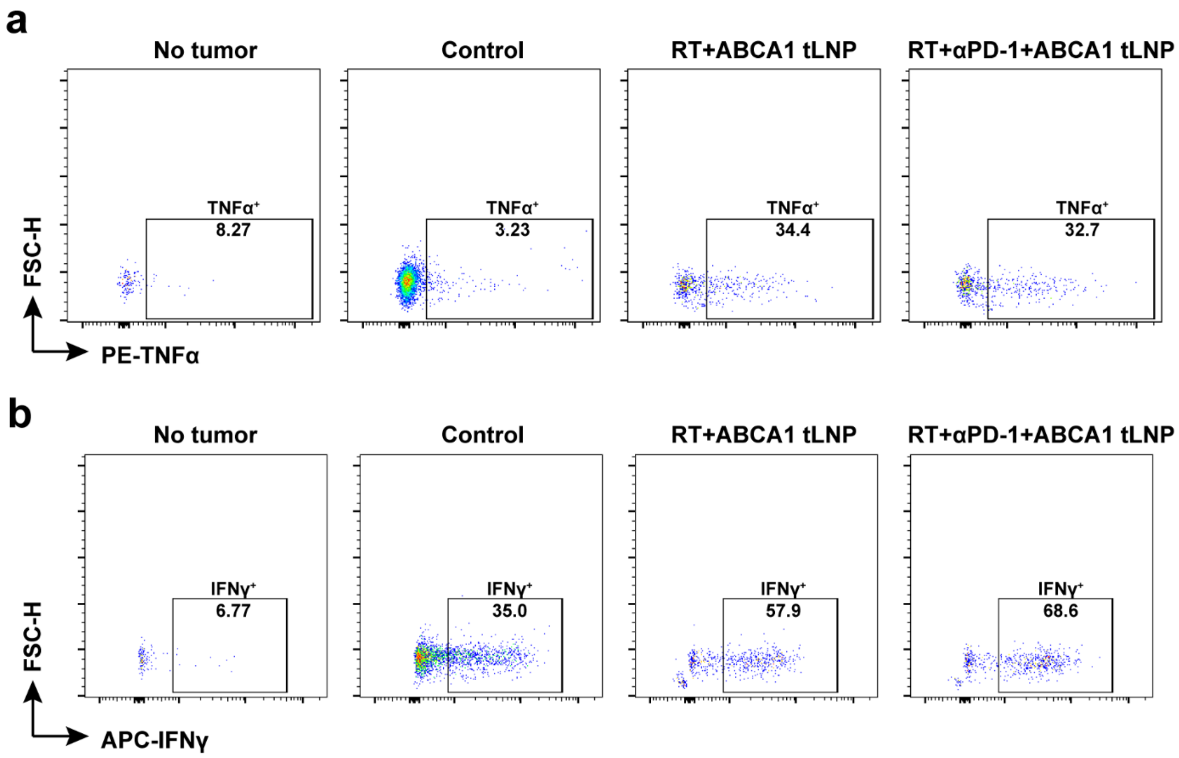


**Figure S5. Flow cytometry plots of TNFα and IFNγ expression in CD8$^+$ T cells. a**, TNFα expression. **b**, IFNγ expression. Treatment conditions included no tumor, control, RT + ABCA1 tLNP, and RT + αPD-1 + ABCA1 tLNP. Representative of n = 4.

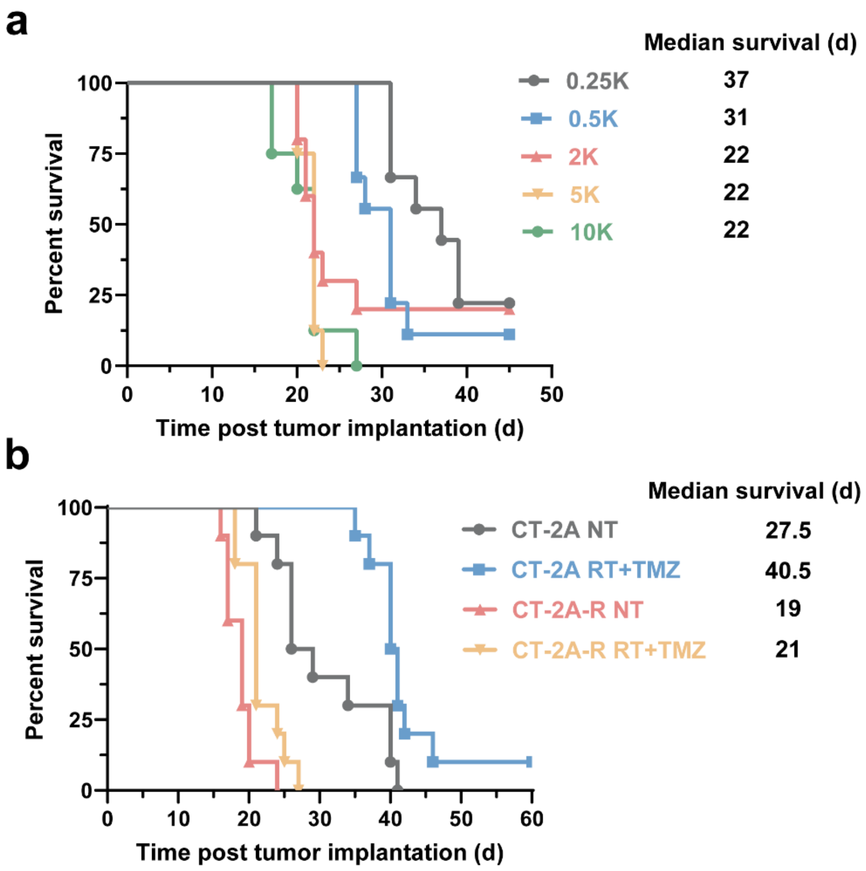


**Figure S6. Kaplan–Meier survival characterization of the CT-2A-R model.** **a**, Kaplan–Meier survival curves of mice implanted intracranially with the indicated numbers of CT-2A-R cells. C57BL/6 mice were intracranially implanted with 10,000, 5,000, 2,000, 500, or 250 CT-2A-R cells per mouse; n = 9. Median survival (days) is indicated. **b**, Kaplan–Meier survival curves comparing parental CT-2A and CT-2A-R tumors under non-treated (NT) or RT + TMZ conditions. n = 10. Median survival (days) is indicated.

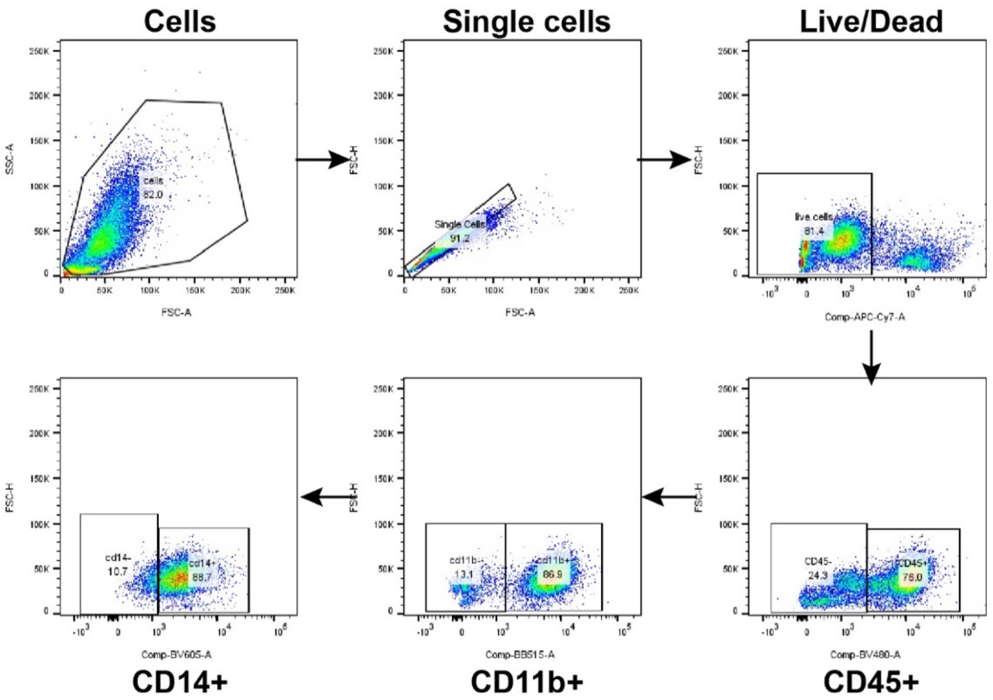


**Figure S7. Flow cytometry gating strategy for human glioblastoma samples.** Cells were sequentially gated on total cells (FSC-A vs SSC-A), singlets (FSC-A vs FSC-H), and live cells. $CD45^+$ immune cells were identified, followed by selection of $CD11b^+$ and $CD14^+$ populations based on marker expression.

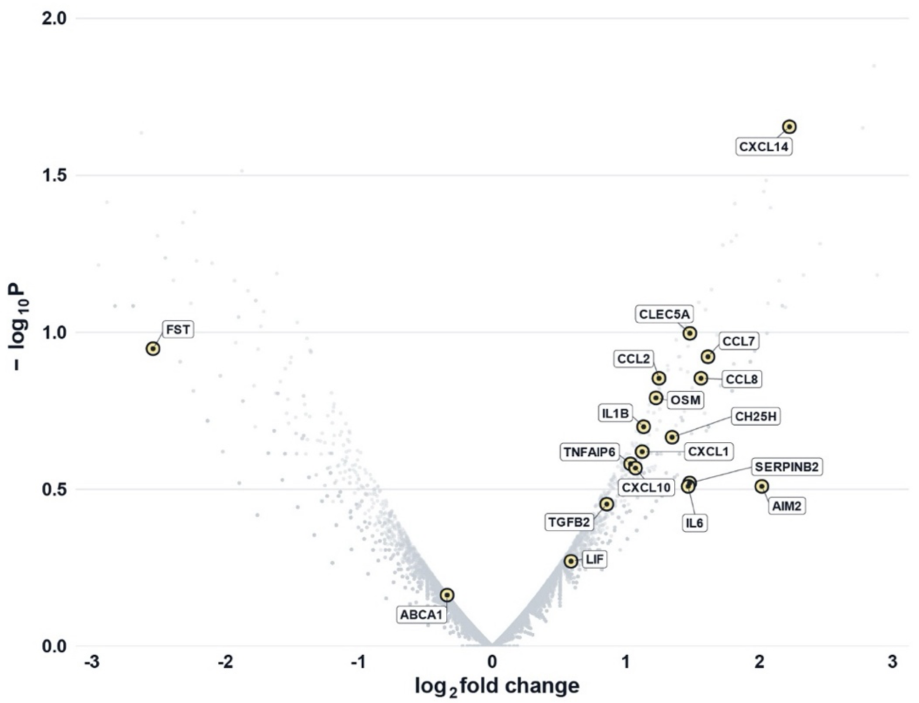


**Figure S8. Volcano plot of differentially expressed genes from bulk RNA-seq comparing ABCA1 tLNP-treated versus control tLNP-treated CD14$^{+}$ myeloid cells from GBM patient sample.** The x-axis indicates $\log_2$ fold change (ABCA1 tLNP versus control tLNP), and the y-axis indicates $-\log_{10}$ (*p* value). Each dot represents a gene; selected representative genes, including chemokines, cytokines, and ABCA1, are highlighted in yellow and labeled. Differentially expressed genes were defined as $|\log_2 \text{fold change}| > 1$ and nominal $p < 0.05$.

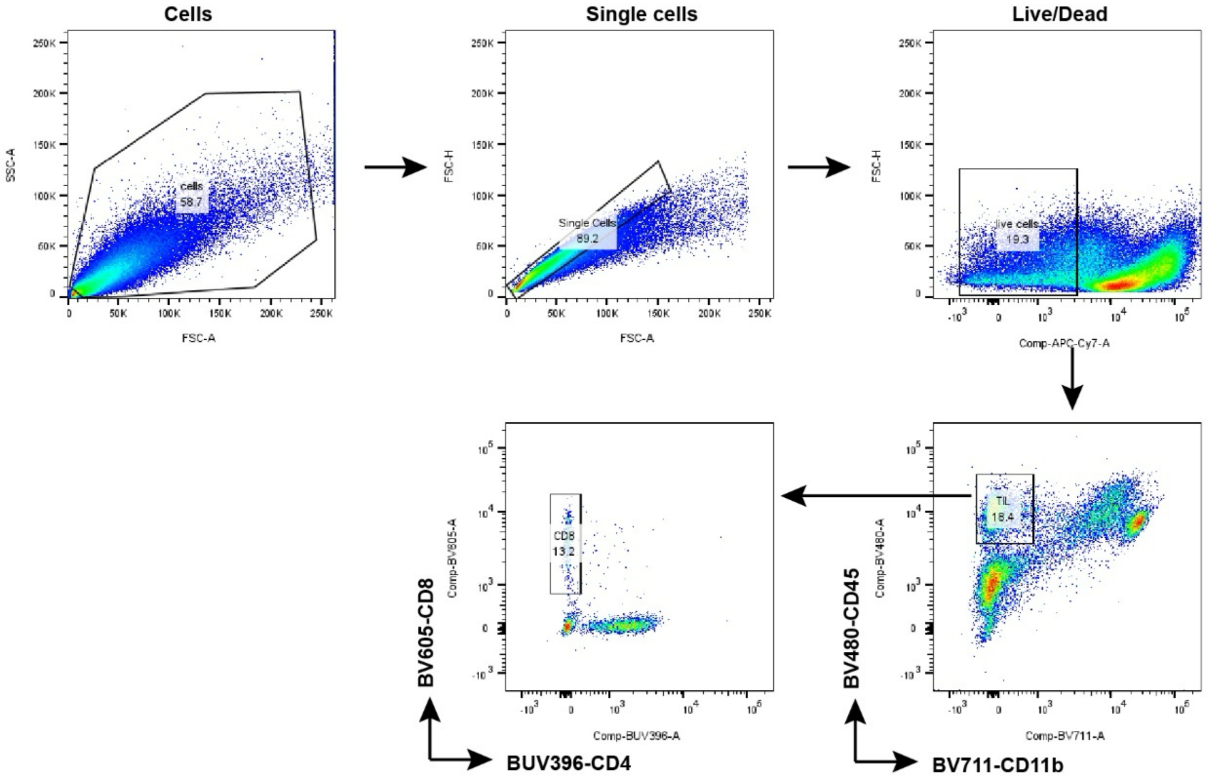


**Figure S9. Flow cytometry gating strategy for Renca tumor samples.** Cells were sequentially gated on total cells (FSC-A vs SSC-A), singlets (FSC-A vs FSC-H), and live cells. $CD45^+$ immune cells were then selected, followed by identification of $CD11b^-$ TILs and $CD8^+$ and $CD4^+$ T cell populations based on marker expression.

**Table S1. Characteristics of GBM patient samples**

| Sample ID | Sex | Race | Ethnicity | WHO grade | IDH status | MGMT Promotor Methylation | Tumor mutation burden (muts/mb) |
|---|---|---|---|---|---|---|---|
| NU03811 | Male | Black | Non-Hispanic | Grade 4 | Wild type | Negative | 6.2 |
| NU03878 | Male | White | Non-Hispanic | Grade 4 | Wild type | Positive | 8.5 |
| NU03922 | Female | White | Non-Hispanic | Grade 4 | Wild type | Positive | 4.6 |
| NU04887 | Female | White | Non-Hispanic | Grade 4 | Wild type | Negative | 6.9 |
| NU04935 | Male | Not reported | Non-Hispanic | Grade 4 | Wild type | Negative | 9.2 |
| NU05064 | Male | White | Non-Hispanic | Grade 3 | Wild type | Negative | 2.3 |

IDH, isocitrate dehydrogenase; MGMT, O6-methylguanine DNA methyltransferase.

**Table S2. siRNA sequences used in this study.**

| siRNA | Sequence (5′→3′) |
|---|---|
| mouse-ABCA1 | GUCCUUACAUCUCAUAGUA[dT][dT] |
| AS/mouse-ABCA1 | UACUAUGAGAUGUAAGGAC[dT][dT] |
| human-ABCA1 | GGUUCUAUGCCCGCUUGAA[dT][dT] |
| AS/human-ABCA1 | UUCAAGCGGGCAUAGAACC[dT][dT] |

AS, antisense strand; [dT], 2′-deoxythymidine 3′ overhang. Sense and antisense strands were annealed to form the duplex siRNA targeting mouse or human ABCA1.

**Table S3. qPCR primer sequences used in this study.**

| Name | Sequence (5′→3′) |
|---|---|
| Abca1-Forward | AAAACCGCAGACATCCTTCAG |
| Abca1-Reverse | CATACCGAAACTCGTTCACCC |
| β-Actin-Forward | CTAAGGCCAACCGTGAAAA |
| β-Actin-Reverse | ACCAGAGGCATACAGGGACA |
| Cxcl9-Forward | CGGACTTCACTCCAACACAGT |
| Cxcl9-Reverse | TTCCTTATCACTAGGGTTCCTCG |
| Cxcl11-Forward | GGCTTCCTTATGTTCAAACAGGG |
| Cxcl11-Reverse | GCCGTTACTCGGGTAAATTACA |

All primers are shown 5′→3′. β-Actin was used as the endogenous reference gene for normalization.

**Table S4. Antibodies used for flow cytometry and immunophenotyping.**

| Antibodies | Clone | Catalog number | Supplier name |
|---|---|---|---|
| Purified anti-mouse CD16/32 | 93 | 101302 | BioLegend |
| BV510 anti-mouse CD45 | 30-F11 | 103138 | BioLegend |
| BV711 anti-mouse/human CD11b | M1/70 | 101242 | BioLegend |
| BV605 anti-mouse CD80 | 16-10A1 | 104729 | BioLegend |
| Alexa Fluor 700 anti-mouse CD86 | GL-1 | 105024 | BioLegend |
| FITC anti-mouse CD206 (MMR) | C068C2 | 141704 | BioLegend |
| BV605 anti-mouse CD8a | 53-6.7 | 100744 | BioLegend |
| PE anti-mouse CD8a | 53-6.7 | 100708 | BioLegend |
| Brilliant Violet 711™ anti-mouse CD4 | RM4-5 | 100557 | BioLegend |
| APC anti-mouse CD223 (LAG-3) | C9B7W | 125210 | BioLegend |
| PE anti-mouse TNF-α | MP6-XT22 | 506306 | BioLegend |
| PE/Cyanine7 anti-mouse CD4 | GK1.5 | 100422 | BioLegend |
| FITC anti-mouse/human CD44 | IM7 | 103006 | BioLegend |
| Pacific Blue anti-mouse CD69 | H1.2F3 | 104524 | BioLegend |
| Alexa Fluor 700 anti-mouse CD25 | PC61 | 102024 | BioLegend |
| PE anti-mouse CD279 (PD-1) | 29F.1A12 | 135206 | BioLegend |
| Alexa Fluor 700 anti-mouse IFN-γ | XMG1.2 | 505824 | BioLegend |
| PE anti-mouse H-2Kb bound to SIINFEKL | 25-D1.16 | 141604 | BioLegend |
| BV510 anti-human CD45 | HI30 | 304036 | BioLegend |
| Spark UV395™ anti-mouse/human CD11b | M1/70 | 101240 | BioLegend |
| BV605 anti-human CD14 | M5E2 | 301834 | BioLegend |
| PerCP/Cyanine5.5 anti-human HLA-DR | L243 | 307630 | BioLegend |
| BV421 anti-human P2RY12 | S16001E | 392106 | BioLegend |
| APC anti-human CD33 | WM53 | 303408 | BioLegend |
| Alexa Fluor 700 anti-human CD8a | RPA-T8 | 301028 | BioLegend |
| PE/Cyanine7 anti-human CD274 (PD-L1) | 29E.2A3 | 329718 | BioLegend |
| BV711 anti-human HLA-DR | L243 | 307644 | BioLegend |

| Antibodies | Clone | Catalog number | Supplier name |
|---|---|---|---|
| PerCP/Cyanine5.5 anti-human CD4 | RPA-T4 | 300530 | BioLegend |
| FITC anti-human CD11b | ICRF44 | 301330 | BioLegend |
| BV711 anti-human CD4 | RPA-T4 | 300558 | BioLegend |
| PE anti-human HLA-A,B,C | W6/32 | 311406 | BioLegend |
